# Hypoxic tumors are sensitive to FLASH radiotherapy

**DOI:** 10.1101/2022.11.27.518083

**Authors:** Ron J. Leavitt, Aymeric Almeida, Veljko Grilj, Pierre Montay-Gruel, Céline Godfroid, Benoit Petit, Claude Bailat, Charles L. Limoli, Marie-Catherine Vozenin

## Abstract

Tumor hypoxia is a major cause of resistance to cancer treatments and especially to radiotherapy (RT) and we wanted to assess whether ultra-high dose rate FLASH RT could overcome this resistance. We engrafted tumor cells of various origins subcutaneously in mice to provide a reliable and rigorous way to modulate oxygen supply via vascular clamping or carbogen breathing. We irradiated tumors using a single 20 Gy fraction at either conventional (CONV) or FLASH dose-rate. Using multiple different subcutaneous tumor models, and in contrast CONV-RT, FLASH-RT retained anti-tumor efficacy under extreme hypoxia. These findings demonstrate that in addition to normal tissue sparing, FLASH-RT overcomes hypoxia-mediated tumor resistance. Follow-up molecular analysis using RNAseq profiling uncovered FLASH-specific inhibition of cell proliferation and translation as well as metabolic shifts that discriminated FLASH-RT from CONV-RT. These data provide new and specific insights into the mechanism of action and identify novel targets for intervention.

## INTRODUCTION

In the field of oncology, resistance to treatment is a major cause of therapeutic failure and therefore overcoming treatment resistance is critical in efforts to enhance the therapeutic index. Resistance to anti-cancer therapies has been investigated intensively and is described as multi-faceted and complex mechanisms that include tumor burden and growth kinetics, tumor heterogeneity, immune suppression, changes in the tumor microenvironment, “undruggable” genomic targets, physical barriers (i.e. blood brain barrier), and hypoxia (reviewed in (1)).

Tumor hypoxia is a primary factor of resistance to radiotherapy (RT), chemotherapy (CT), and immunotherapy (IT). Reduced oxygen tension is very common in solid tumors (2–4), as it develops in response to a dense and metabolically active tumor-cell population associated with poor, uneven, and morphologically abnormal vasculature resulting in acute (proximal to host blood vessel) or chronic (distal to any vasculature) hypoxia. While the lack of oxygen itself is the primary cause of radioresistance, irregularities in the vascular network are mainly responsible for CT and IT resistance preventing drug access to the tumor. At the cellular level, hypoxia causes rapid activation of hypoxia-inducible factor (HIF) transcription factors (5) as acute survival mechanisms. HIFs transactivate an array of genes involved in angiogenesis, pH balance, cell apoptosis, and a shift to anaerobic metabolism essential for tumor survival and resistance to treatment. Multiple attempts to overcome tumor hypoxia to facilitate effective treatment have been proposed including hypoxia-activated prodrugs, hyperbaric oxygen breathing, 60% supplemental oxygen, allosteric hemoglobin modifiers, molecules that improve oxygen diffusion, and oxygen transport agents (hemoglobin-based or fluorocarbon-based) (6). Most of these strategies have led to negative results on tumor control and enhanced normal tissue toxicity. To date, no effective anti-cancer therapy against hypoxic tumors are available.

Previously, our team developed a novel radiotherapy approach called FLASH-RT based on ultra-high-dose-rate (UHDR) irradiation beams (7–13). The main interest of FLASH-RT is its capacity to enhance the therapeutic window by sparing normal tissue from radiation-induced toxicities while eliciting the same tumor kill when compared to iso-doses of conventional dose-rate RT, a biological effect that has been called the “FLASH effect”. While currently the availability of FLASH-capable clinical beam remains the main obstacle for the clinical translation of FLASH-RT (14), understanding the differential impact of FLASH-RT has at the normal tissue *versus* tumor level remains an important goal to further decipher the mechanisms of the FLASH effect (15) and facilitate clinical translation. Classical radiobiological dogma posits that radiochemical oxygen depletion induced at UHDR would cause a transient protective hypoxia in the healthy tissue, while hypoxic tumors would be less affected (16). This hypothesis has now been challenged by experimental measurements, where radiochemical oxygen depletion following UHDR in normal tissues and tumors was found to be minimal (< 0.2%) after exposure to therapeutic doses (2-10 Gy) (17). In another hypothesis, FLASH-RT has been proposed to protect stem cells known to be preferentially located in hypoxic niches (18). However, this also suggests that hypoxic tumors and cancer stem cells would be protected, contrary to the dose-rate independence of tumor cell kill observed in FLASH-RT studies. Clearly, further in-depth investigations are needed to evaluate the impact of FLASH in hypoxic tumors.

In light of the foregoing, it is noteworthy that nearly every preclinical cancer model tested has shown identical tumor response when FLASH-RT and conventional-dose-rate RT (CONV-RT) were compared (7,17,19–22). Importantly, to date, there has not been any investigation performed to evaluate the efficacy of FLASH-RT on extremely hypoxic, radioresistant tumors.

To rectify this critical gap in knowledge, we focused the present work on studying the impact of intratumoral oxygen tension on the tumor response to FLASH-RT. Here we implemented different validated models of subcutaneous tumors that provide a reliable and rigorous way to modulate intratumoral oxygen tension by vascular clamping or carbogen breathing. Our results show that, contrary to conventional dose rate RT, FLASH-RT maintains its anti-tumor efficacy ***even under extreme hypoxia***. We performed subsequent molecular analysis using RNAseq profiling at acute and relapse time points, with data pointing toward specific inhibition of proliferation and translation-associated genes as well as metabolic shifts following FLASH-*versus* CONV-RT. More specifically, we identified glycolysis as a possible escape/survival pathway following exposure to FLASH-RT. Therefore, we delivered a U.S. Food and Drug Administration (FDA)-approved glycolysis inhibitor (trametinib) as adjuvant treatment after FLASH-RT resulting in significantly delayed time to relapse.

## MATERIALS AND METHODS

### Animal experiments

Female Swiss Nude mice (NU(Ico)-*Foxn1^nu^*; Charles River FR) were purchased for subcutaneous tumor experiments using the U-87 MG, H454, SV2, and mEERL95 tumor models. Female C57BL/6J mice (Charles River FR) and C57BL/6JRJ (Janvier Labs FR) were purchased for subcutaneous tumor experiments using SV2 and mEERL95 tumor models, respectively. Male Swiss Nude mice (NU(Ico)-*Foxn1^nu^*; Charles River FR) were purchased for subcutaneous tumor experiments using the RKO and RKO.AS45.1 tumor models. Animal experiments were approved by the Swiss Ethics Committee for Animal Experimentation (VD 3241 – VD 3603 – VD 3670 – VD 3797) and performed within institutional guidelines.

### *In vivo* tumor models, irradiations and follow-up

The U-87 MG subcutaneous human glioblastoma model consists of injecting 10^7^ U-87 MG human GBM cells in 100 μl PBS in the right flank of female Swiss Nude mice. For SV2 lung adenocarcinoma, 10^6^ cells were injected in 100μl PBS + 40% Corning Matrigel Basement Membrane Matrix (356234, Corning) in the flank of C57BL/6J mice. For mEERL95 head & neck tumors, 5x10^5^ cells were resuspended in 100 μl PBS and injected in the flank of C57BL/6JRJ mice. Irradiations were performed when tumor volume reached a mean between 60 - 80 mm^3^. Mice were randomly assigned to one of three different irradiation groups (NIR, CONV-RT, and FLASH-RT) and to one of three different oxic-conditions groups (hypoxia, physioxia, hyperoxia). Initial mean tumor volumes were similar in all groups.

Tumors were irradiated by positioning the tumor-bearing part of the skin in extension behind and in contact with the opening of the 1.7 cm diameter graphite applicator, in order to limit the dose to the intestines. A 5 mm solid water plate was placed behind the skin to ensure homogenous dose delivery. For irradiations under hyperoxic conditions, mice were anesthetized with isoflurane and carbogen (95% O_2_, 5% CO_2_) for at least 20 min, including the irradiation time. For irradiations in hypoxic conditions, tumors were clamped with a vascular clamp at least 15 min before and during the irradiation. All tumors were treated with a single dose of 20 Gy. For all regimen, FLASH and CONV irradiation modalities were compared.

### Intratumoral oxygen tension measurements using the OxyLED system and Oxyphor PtG4 probe

On the day of the experiment, animals were anesthetized as described in the supplemental methods. Once anesthetized, 25uL of Oxyphor PtG4 probe (200 μM; Oxygen Enterprises Ltd., USA) was injected intravenously. After 30 minutes to ensure the spread and accumulation of the probe, the OxyLED optical fiber (Oxygen Enterprises Ltd., USA) was placed about 1 cm close from the tumor to perform the measurement of the intratumoral oxygen tension. All measurements were performed under anesthesia. Measurements were taken for all 3 oxygenation conditions, n=3 / condition.

### Adjuvant trametinib and vehicle treatments

Trametinib (C3822, ALSACHIM) was dissolved in DMSO (D4540, Sigma-Aldrich) to a concentration of 10 mM (∼6.15 mg/mL) and kept in aliquots at −80°C until use. On treatment day, an aliquot of working solution was thawed and added to sterile PBS for gavage. DMSO was added to PBS for the vehicle treatment. Trametinib was administered daily over the course of 2 weeks (days 1 – 15 post-RT) at a concentration of 1 mg/kg body weight by oral gavage in 200 μl total volume.

### Statistical analyses

Statistical analyses were carried out using GraphPad Prism (v9.1) software. Results were expressed as mean values + SEM, and all analyses considered a value of P ≤ 0.05 to be statistically significant unless specified otherwise.

### Supplemental Methods

Supplemental Methods include experimental details on the irradiation device and parameters, cell culture protocols, tumor volume calculations, animal anesthesia, Pimonidazole hypoxia confirmation, RNA extraction and High-Throughput Sequencing, Bioinformatic Analysis, and selected statistical analyses.

### Data and materials availability

The data discussed in this publication have been deposited in NCBI’s Gene Expression Omnibus (43) and are accessible through GEO Series accession number GSE223607 (https://www.ncbi.nlm.nih.gov/geo/query/acc.cgi?acc=GSE223607). All data and code are available upon request

## RESULTS

### Severe hypoxia induces resistance to CONV but not FLASH-RT in various experimental subcutaneous tumor models

To determine the impact of intratumoral oxygen tension on tumor response following a single dose of FLASH-RT, we used a human glioblastoma (GBM; U-87 MG) xenograft model implanted subcutaneously in the flank of immunodeficient female Swiss Nude mice. We modulated the tumor oxygen tension using a vascular clamp (hypoxia) or carbogen breathing (hyperoxia). Results showed that a single FLASH-RT dose of 20 Gy was more effective to delay the growth of hypoxic tumors compared to a similar dose of CONV-RT (Fig. 1A). We found no statistical difference in tumor growth between hypoxic and physioxic tumors following FLASH-RT and animal survival data (based on time to interruption criteria using Kaplan–Meier curves) were similar in the physioxia groups after both FLASH-and CONV-RT and in the hypoxia group after FLASH-RT (Fig. 1B). The mean doubling time of U-87 tumors was about 20 days (95% CI from 18.4 to 24.1 days) when non-irradiated (NIR) and unmodified in oxygen tension (Fig. 1C). After CONV-RT, the mean doubling time of the tumor was 33.7 days in physioxia, 26.6 in hypoxia and 40.7 days in hyperoxia. After FLASH-RT, the mean doubling time of the tumor was 31.5 days in physioxia, 30.6 in hypoxia and 42.2 days in hyperoxia. There was no significant difference between CONV-and FLASH-RT mean doubling times in physioxia (P = 0.175) or hyperoxia (adjusted p-value = 0.646), whereas in hypoxic condition FLASH-RT remained efficacious to delay tumor growth (P = 0.519 vs FLASH-physioxia and P = 0.0148 vs CONV-hypoxia). To confirm the different oxic conditions, we measured intratumoral oxygen tension before treatment via real-time oxygen readings using the OxyLED detector and Oxyphor PtG4 probe (Oxygen Enterprises Ltd., Fig. 1D) and hypoxia was confirmed post-treatment using pimonidazole staining (Fig. S1).

**Fig. 1.**
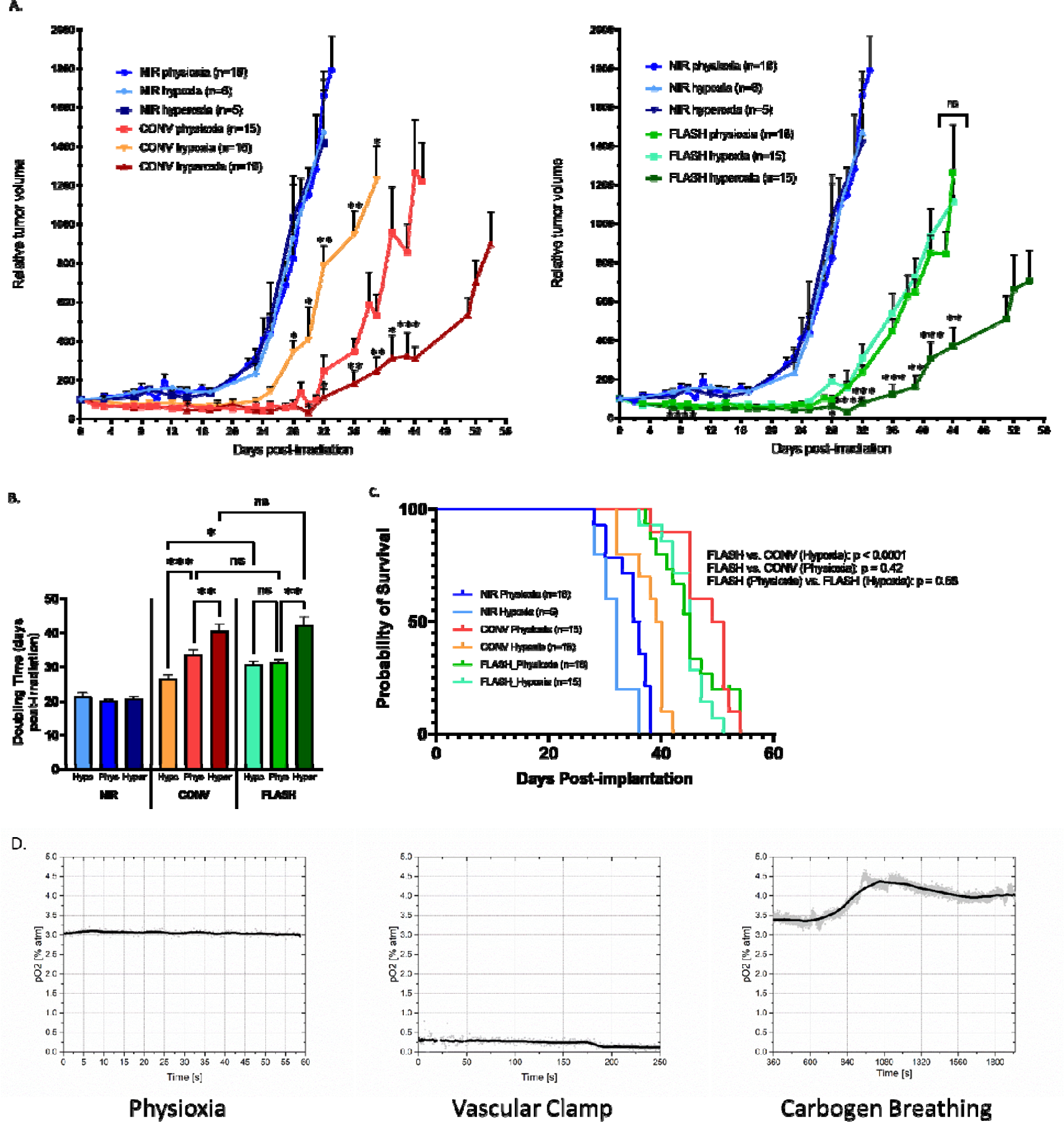
FLASH-RT, unlike CONV-RT, maintains equivalent tumor control even in severely hypoxic tumors. (A) Relative tumor volume of U-87 MG implanted subcutaneously in the flank of female Nude mice treated with 20 Gy single fraction delivered with CONV (left) or FLASH (right) radiation therapy with different oxygenation conditions (physioxia – hypoxia – hyperoxia). Mean relative tumor volume + SEM, N = 5-18 animals per group. P values derived from multiple row-wise Mann-Whitney test against the physioxia group: *P < 0.05; **P < 0.01; ***P < 0.001; ****P < 0.0001; ns, not significant. (B) Doubling times calculated for each group. Mean days post-RT + SEM, N=5-18 animals per group. P values derived from unpaired two-sided t test with Welch’s correction: *P < 0.05; **P<0.01; ***P < 0.001; ns, not significant. Kaplan–Meier survival curves (C) for animals stratified by radiation modality and tumor oxygenation. P values derived from log rank (Mantel-Cox) test. (D) Representative real-time oxygen measurements from a subcutaneous U-87 MG tumor in vivo taken using Oxyled/OxyPhor PtG4 (Oxygen Enterprises Ltd.) phosphorescent probe with platinum core injected intravenously and detected using the laser attachment and detector are shown for the three different oxygenation conditions (physioxia, vascular clamp, and carbogen breathing). Signal averaging line shown in bold.

We were able to replicate the enhanced efficacy of FLASH-RT against hypoxic tumors compared to CONV-RT multiple times by different experimenters in the same model (Fig. S2) and further confirm this finding using H454 mouse GBM cells grafted in the Swiss nude mice (Fig. S3). Additionally, we grafted SV2 mouse lung cancer and mEERL95 mouse head-and-neck cancer cell lines in both Swiss Nude mice and immunocompetent C57BL/6J mice yielding similar results (Figs. S4-S5), indicating a minimal contribution of the adaptive immune system to the antitumor effect of FLASH-RT in hypoxic tumors. These data show an enhanced efficacy of FLASH-RT in hypoxic tumors as compared with CONV-RT.

### Existence of a FLASH-RT-specific genomic imprint

To determine the mechanism of the anti-tumor efficacy triggered by FLASH vs CONV-RT, we performed bulk RNAseq studies at early (24 hours post-RT) and more protracted (one-week post-tumor-recurrence) time points. The analysis included only the tumor component, as subcutaneously-grown tumor nodules are mainly composed of tumor cells. Additionally, following alignment, we applied a de-convolution step to discriminate between human component (tumor) and mouse tissue contamination (host). Overall, we observed more top-level transcriptional differences in the 24hour time point (Fig. 2A, PC (principal component) 1 accounting for 78% of variance). As shown on the heatmap (Fig. 2B), FLASH-RT samples (green) tended to cluster further away from NIR samples (blue) than CONV-RT (red). For the samples taken at later time point post-RT, PCA (principal components analysis; Fig. 2C, PC1 accounting for 39% of variance) revealed a good clustering of the NIR groups (circle, left), whereas all of the RT-treated tumors clustered together without distinction based on RT modality or oxygenation condition. We confirmed this pattern with the heatmap (Fig. 2D).

**Fig. 2.**
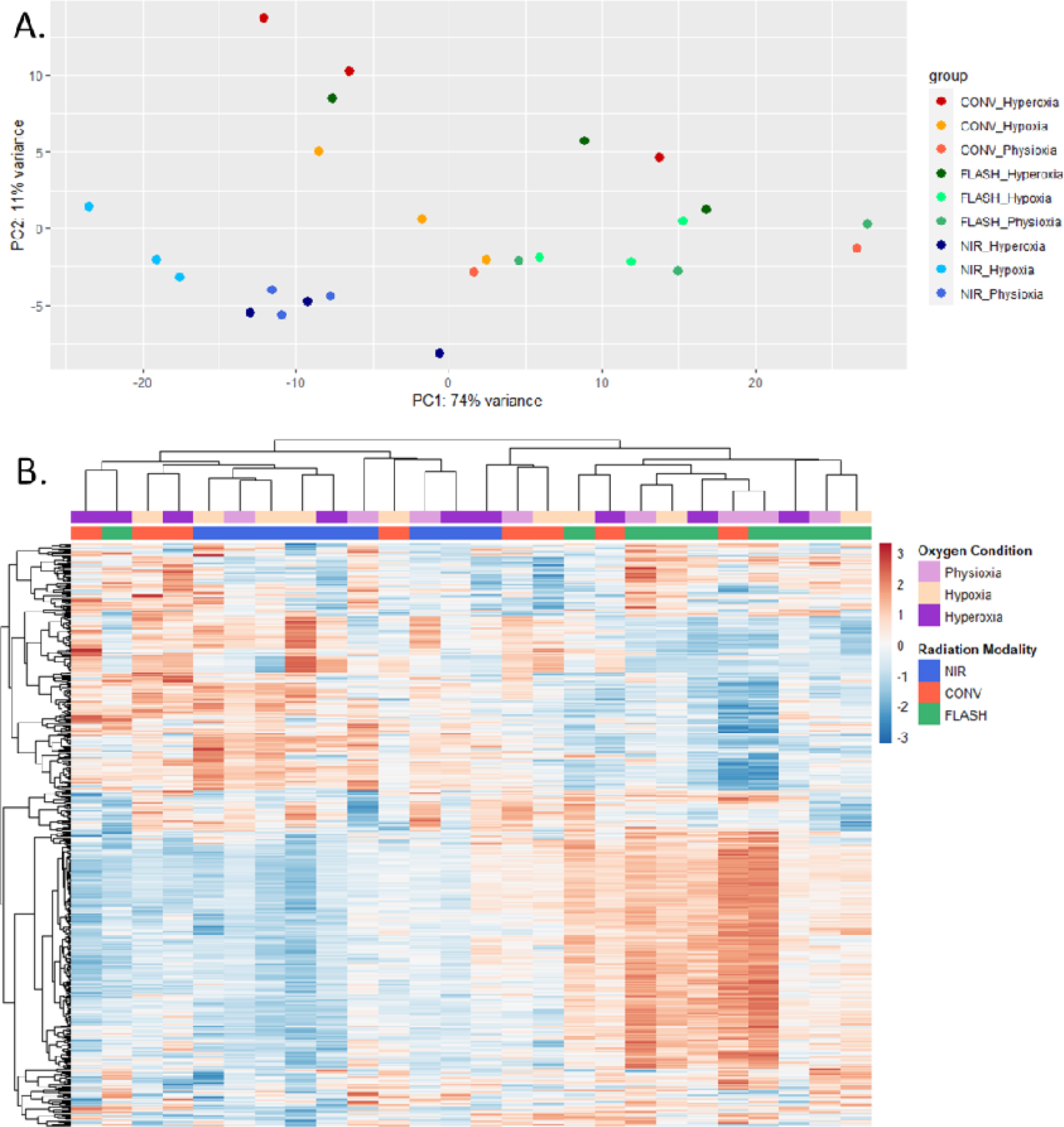
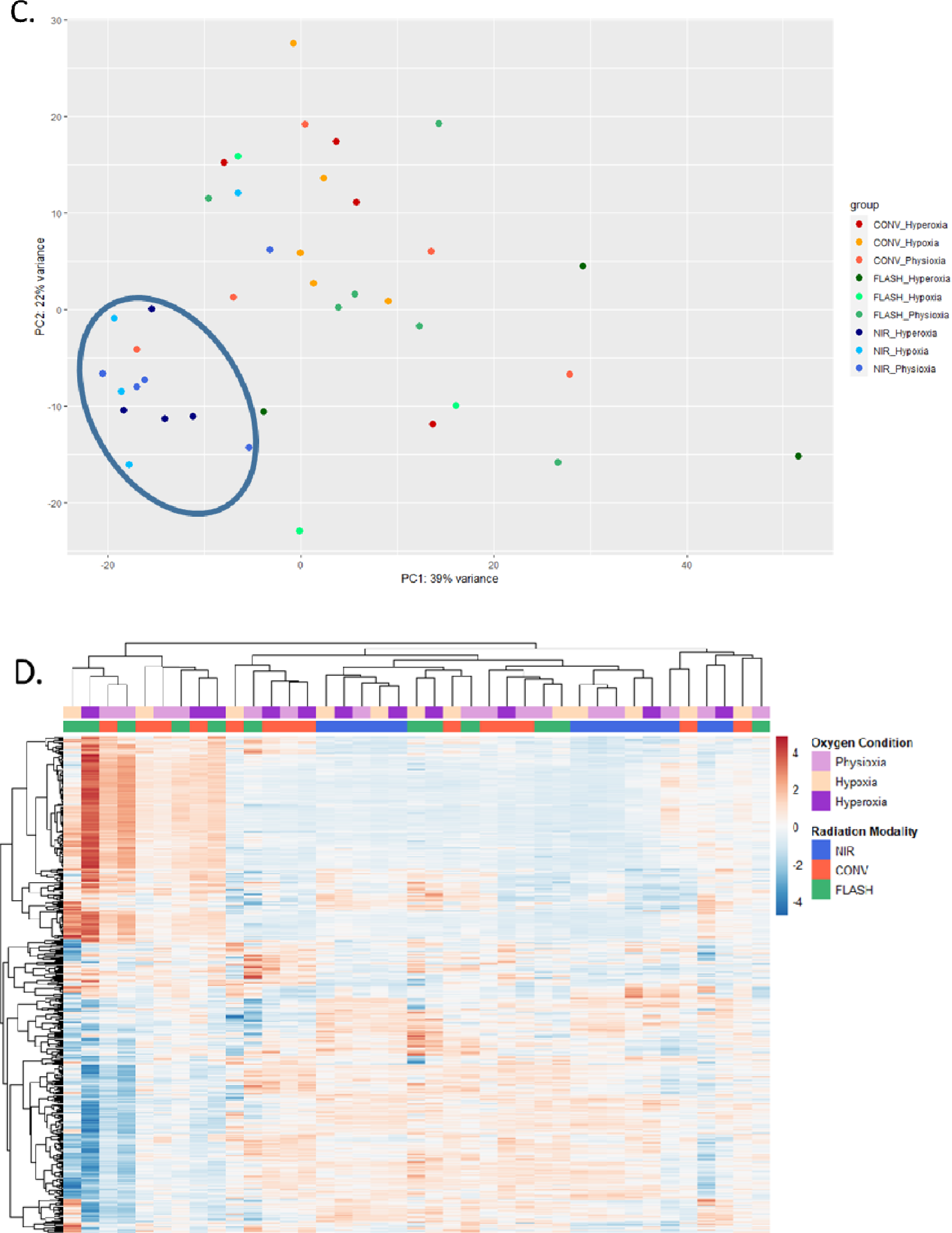
RNAseq Data Overview. Principal components analysis (PCA) to visualize sample-to-sample distances based on the top two principal components (PC1 and PC2) for 24 hour (A) and late (C) time points. Circle (blue) to highlight the cluster of non-irradiated controls at the late time point. Heatmaps for both time points (B and D, respectively) for the top 500 genes – columns (samples) and rows (genes, Z-score) clustered hierarchically.

### FLASH-RT efficacy in hypoxic conditions is associated to a specific genomic imprint

Focusing on the physioxic and hypoxic conditions 24 hours post-RT, the PCA and heatmap showed that treatment modalities and tumor oxygen tensions associated with the longest tumor growth delay (CONV physioxia, FLASH physioxia, and FLASH hypoxia) are clustering together (Figs. 3A and 3B, right).

**Fig. 3.**
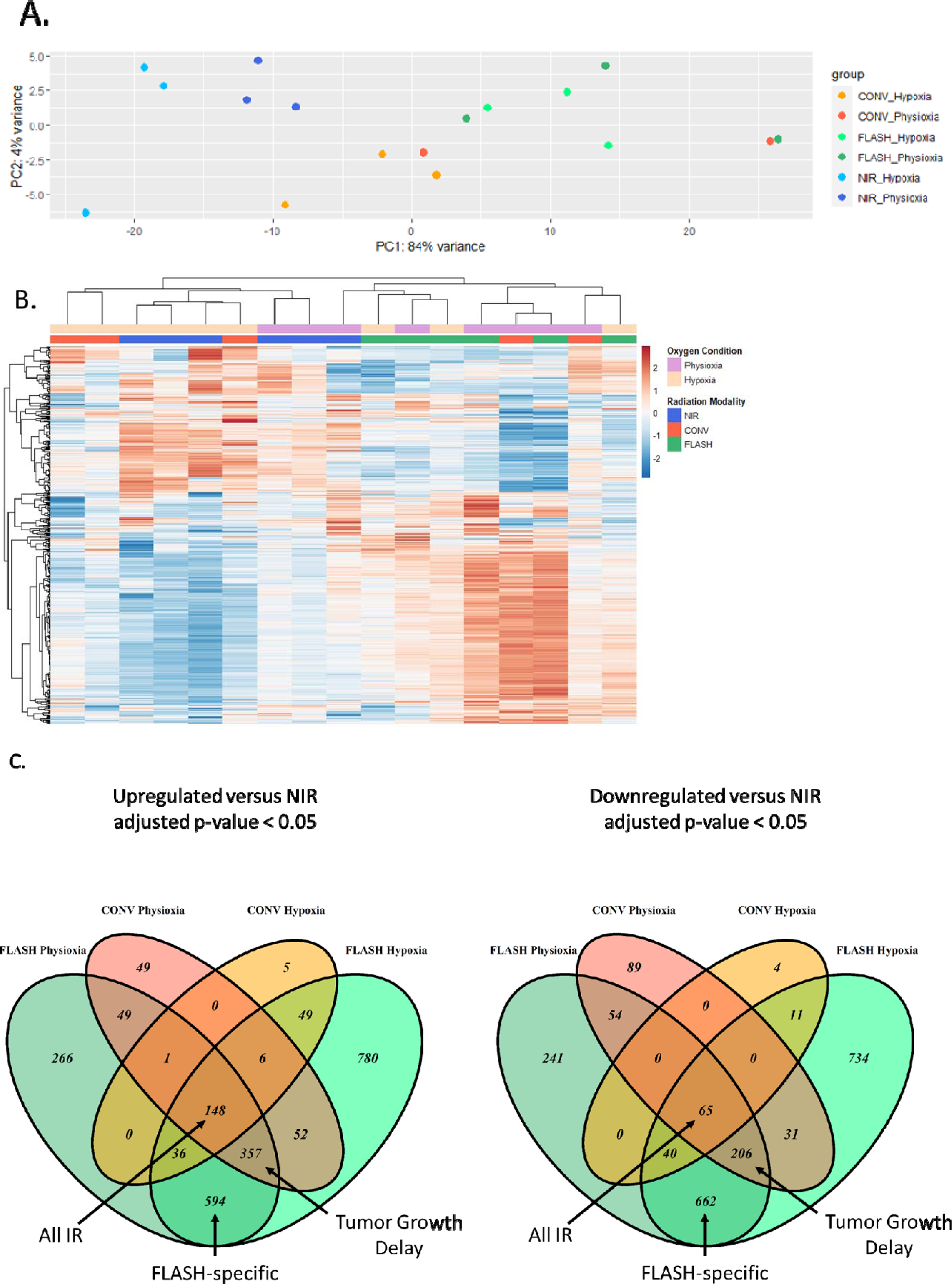
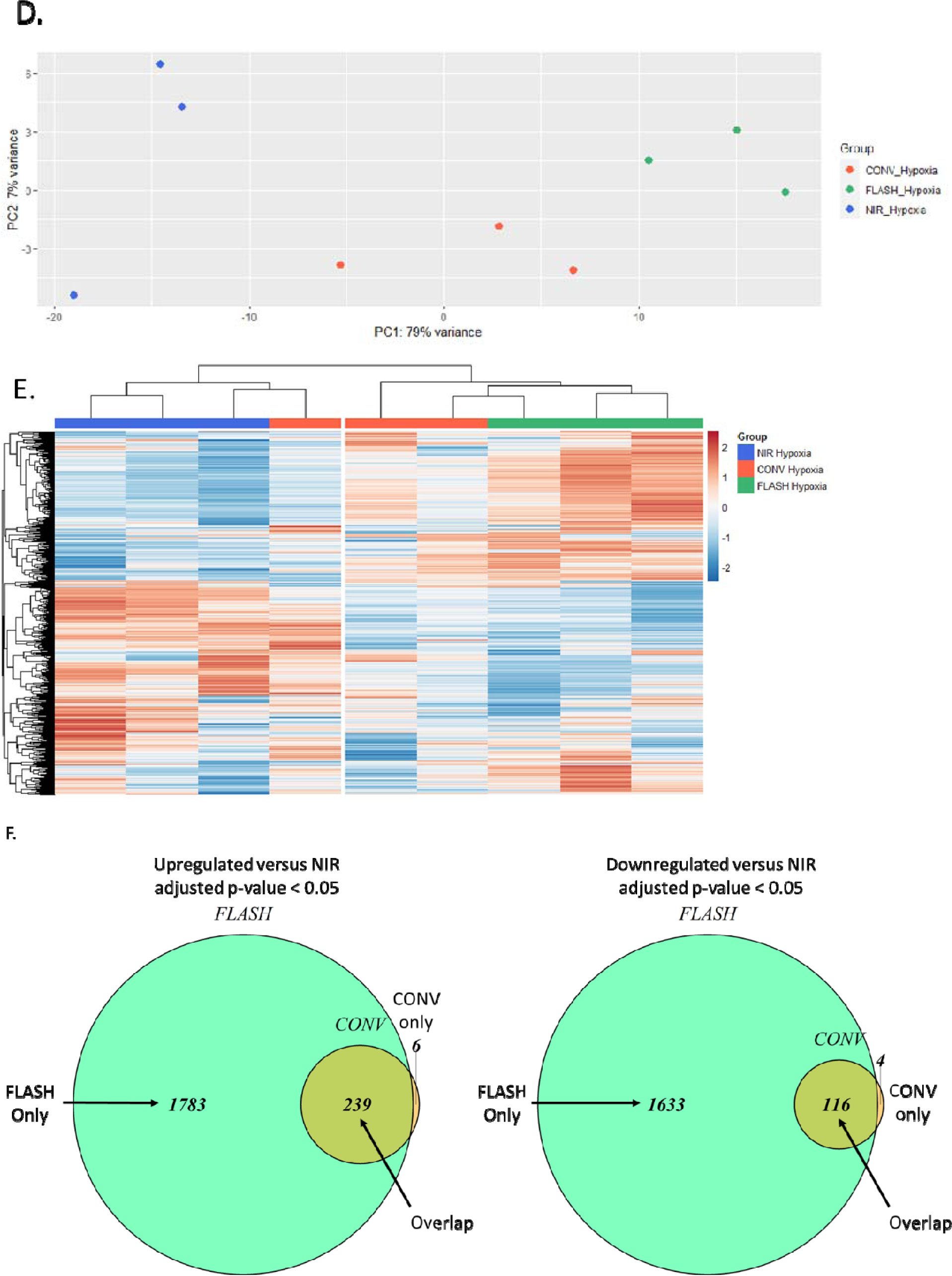
Tumor Growth Delay and FLASH-RT-specific clusters identified both for physioxic and hypoxic conditions. (A) Principal components analysis (PCA) to visualize sample-to-sample distances based on the top two principal components (PC1 and PC2) for the physioxia and hypoxia conditions. (B) Heatmap with the same samples showing the top 500 genes – columns (samples) and rows (genes, Z-score) clustered hierarchically. (C) Venn diagrams depicting the overlap of genes significantly (adjusted p-value < 0.05 from DESeq2 output) upregulated or downregulated in treated groups versus non-irradiated controls. (D) PCA to visualize sample-to-sample distances based on the top two principal components (PC1 and PC2) for only the hypoxia condition. (E) Heatmap with the hypoxia samples showing the top 500 genes – columns (samples) and rows (genes, Z-score) clustered hierarchically. (F) Venn diagrams depicting the overlap of genes significantly (adjusted p-value < 0.05 from DESeq2 output) upregulated or downregulated in treated groups versus non-irradiated controls.

Using the DESeq2 output, we made lists of all differentially expressed genes in irradiated groups *versus* the NIR control group and determined all overlaps. We identified three main overlap profiles. **FLASH-specific**: genes changed significantly *versus* NIR in both FLASH physioxia and FLASH hypoxia groups, but not in neither of the CONV-RT groups with 594 upregulated genes and 662 downregulated genes found; **Tumor Growth Delay**: genes changed significantly in FLASH hypoxia, FLASH physioxia and CONV physioxia groups *versus* NIR, but not the CONV hypoxia group with 357 upregulated and 206 downregulated genes found; and **All-IR**: genes changed significantly in all irradiated groups *versus* non-irradiated control with 148 upregulated and 63 downregulated genes found (Fig. 3C).

Applying the same analysis exclusively for hypoxic conditions 24 hours post-RT, the PCA showed distinct clusters for CONV hypoxia, FLASH hypoxia, and NIR hypoxia groups (Fig. 3D), clearly confirmed by the heatmap (Fig. 3E). This analysis resulted in only three possible profiles: genes altered in FLASH or CONV-RT samples only, and genes altered in both FLASH and CONV-RT samples. Using the same overlap strategy, we found that the FLASH-only signature in the hypoxic condition was associated with 1783 genes uniquely upregulated and 1633 genes uniquely downregulated (Fig. 3F). Using all overlap groups (FLASH-specific, Tumor Growth Delay, All IR, FLASH only, Overlap, and CONV only) we performed an overrepresentation analysis (ORA) of multiple databases including Wikipathways, Kyoto Encyclopedia of Genes and Genomes (KEGG), and Gene Ontology (GO). Based on a global survey of the enrichments, we selected three main altered clusters of enrichments: 1) cell cycle, 2) translation and ribosome, and 3) HIF1 signaling and metabolism.

### FLASH is more efficient than CONV-RT at inhibiting cell cycle

We performed ORA analyses of downregulated genes, which showed enrichment for cell cycle-related pathways for the Wikipathways and KEGG databases in physioxic and hypoxic conditions (Fig. 4 A-B). While we found an enrichment of cell cycle and DNA replication in the genes downregulated in the All IR overlap, we detected further significant enrichments in the Tumor Growth Delay overlap, and even more in the FLASH-specific overlap, suggesting an enhanced FLASH-induced inhibition of cell-cycle-related pathways. Similarly, in analyzing the downregulated genes in only the hypoxia condition, we found enrichment for cell-cycle-related pathways (Fig. 4 C-D) in both FLASH-RT only and Overlap groups and just minor enrichment in CONV Only. This observation supports the trend of expanded downregulation of cell cycle-related genes by FLASH-RT. By directly comparing the change in gene expression following FLASH and CONV-RT in hypoxia and determining enrichments using gene set enrichment analysis (GSEA), we highlighted the downregulation of cell-cycle related genes as shown in Fig. 4E and GSEA plot (Fig. 4F). For the cell cycle pathway (Fig. 4G), we observed a substantial downregulation (blue) in FLASH hypoxia *vs.* CONV hypoxia, generally for the S (replication), G2 (growth and preparation for division), and M (mitosis) phases. Consistent with cell cycle inhibition, we found that p53 effector GADD45 was upregulated after FLASH hypoxia (middle, red), which could explain the inhibition of the Cyclin B / CDK1 complex formation (of which both transcripts were found to be significantly downregulated).

**Fig. 4.**
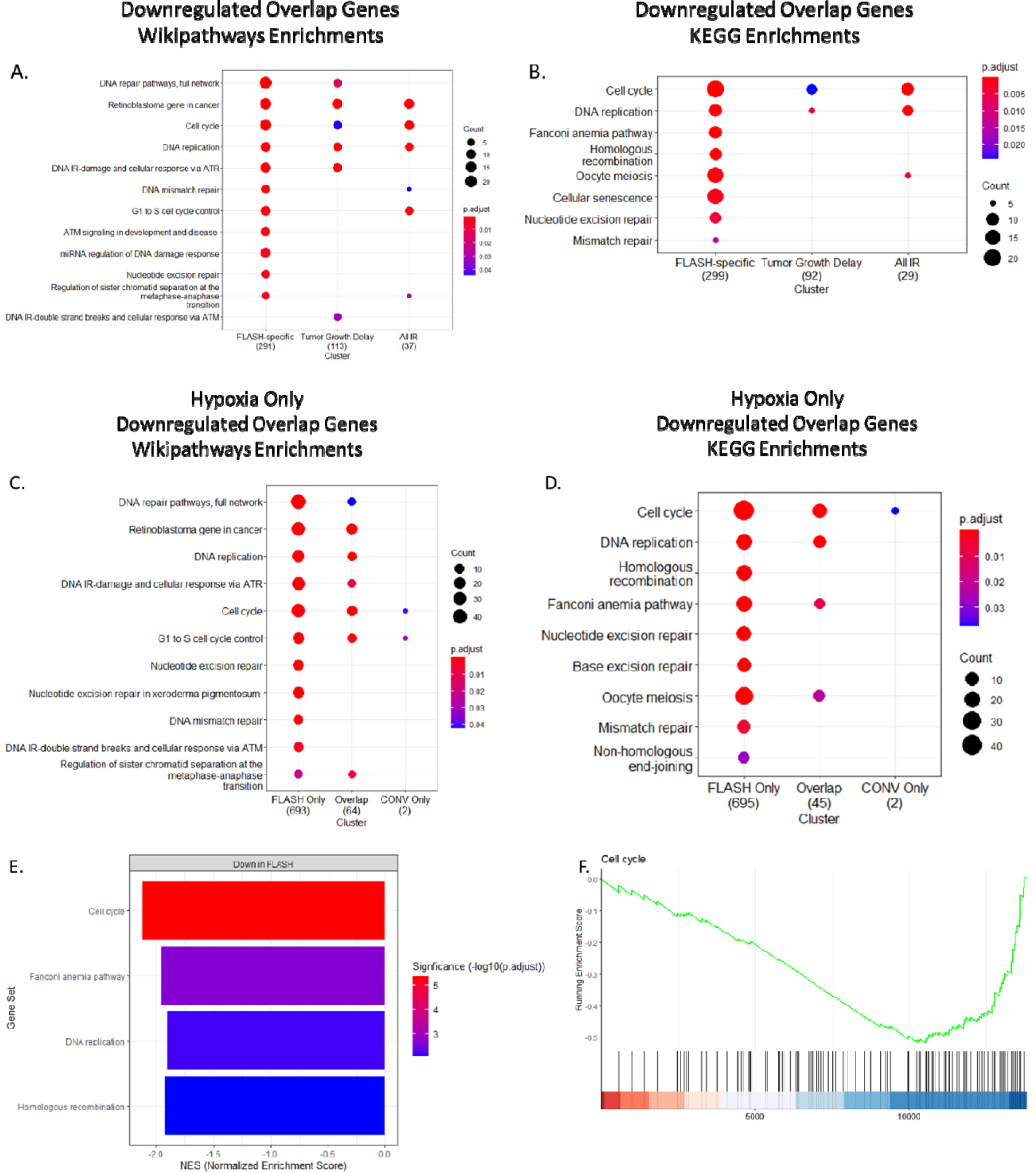
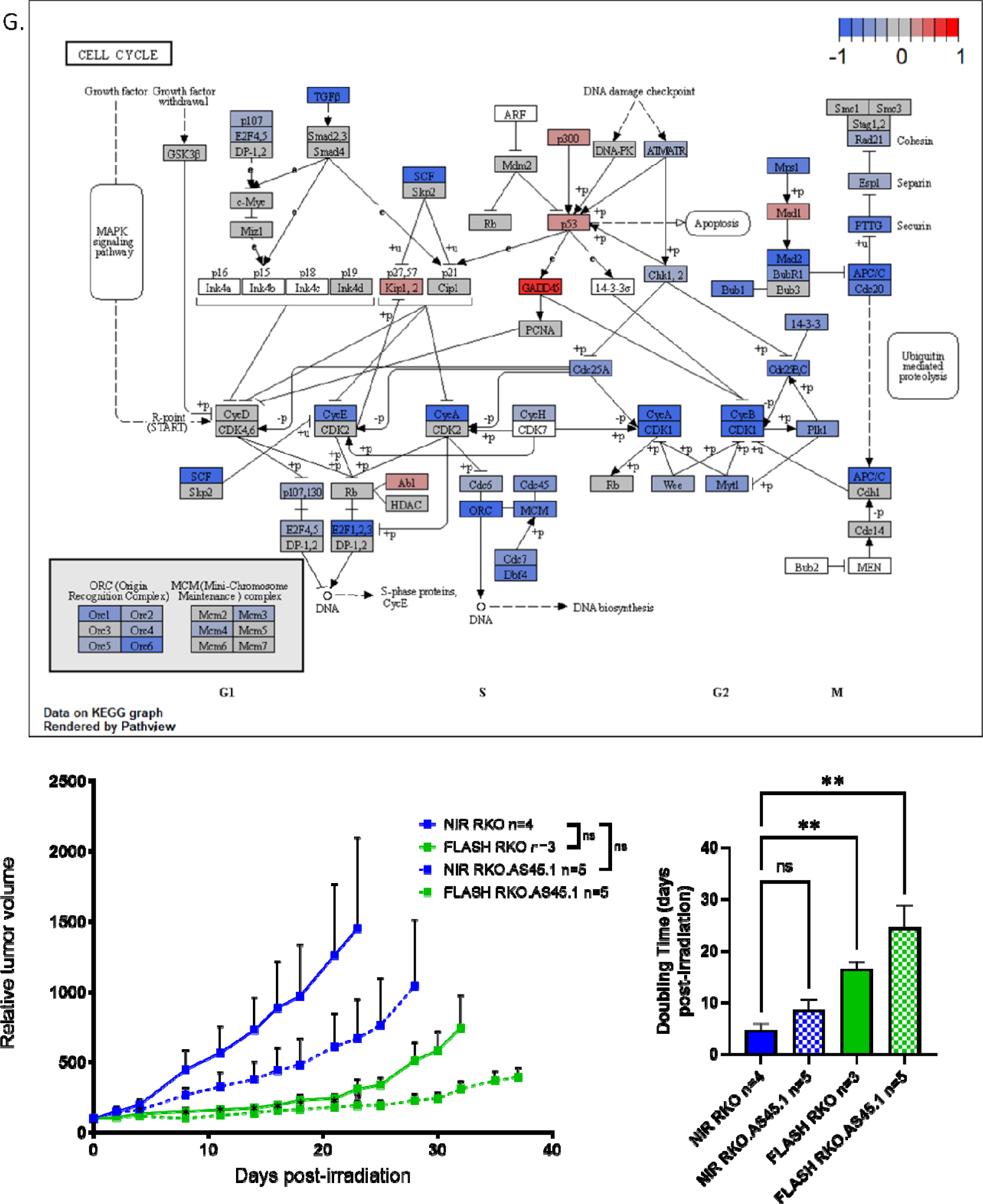
Cell cycle and DNA repair pathways further enriched in significantly downregulated genes in FLASH-RT-treated group, both in hypoxia and physioxia. Overrepresentation analysis (ORA) for pathways from the Wikipathways database for differentially expressed gene (DEG) overlap clusters in physioxia (A) and hypoxia (C). ORA for pathways from the Kyoto Encyclopedia of Genes and Genomes (KEGG) database for DEG overlap clusters in physioxia (B) and hypoxia (D). For the hypoxic condition: select cell cycle-related pathways (E) enriched in FLASH-RT versus CONV-RT using gene set enrichment analysis (GSEA) for the KEGG database. (F) GSEA plot of the negative (down in FLASH-RT) enrichment for the KEGG cell cycle pathway. (G) KEGG cell cycle pathway with overlay of the log2 fold changes of FLASH-RT versus CONV-RT (hypoxia) from the DESeq2 output. (H) Relative tumor volume of human RKO and RKO.AS45.1 colon carcinoma tumors implanted subcutaneously in the flank of male Swiss nude mice untreated or treated with 20 Gy single fraction delivered with FLASH-RT. Mean relative tumor volume + SEM, N = 3-5 animals per group. P values derived from multiple row-wise Mann-Whitney tests against the NIR RKO group, for both tumor models: *P < 0.05; ns, not significant. (I) Doubling times calculated for each group. Mean days post-RT + SEM, N=3-5 animals per group. P values derived from unpaired two-sided t test with Welch’s correction: **P < 0.01; ns, not significant.

We conducted a test to support our findings by examining the sensitivity of GADD45-overexpressing RKO human colon carcinoma xenograft tumors. We observed that the GADD45-overexpressing (RKO.AS45.1) tumor growth was delayed compared to the parental (RKO) tumor growth in the absence of irradiation, although the difference was not statistically significant (Fig. 4H). After administering a single 20 Gy dose of FLASH-RT, we found that the RKO.AS45.1 tumor growth was significantly delayed compared to the RKO NIR tumor growth, while the RKO tumor growth did not show any statistically significant differences at any time point. We calculated the mean doubling time of RKO tumors to be 4.7 days, while GADD45 overexpression alone increased it to 8.6 days (P = 0.147; Fig. 4I). Further, FLASH-RT alone increased the mean doubling time to 16.6 days post-RT (P = 0.0024), and finally, we found that the combination of GADD45 overexpression and FLASH-RT increased it even further to 24.5 days post-RT (P = 0.0077). Altogether, these data demonstrate that we have identified a role for GADD45 and G2/M checkpoint inhibition in enhancing tumor cell sensitivity to FLASH-RT.

### Inhibition of translation and ribosomal biogenesis is FLASH-specific

Using the same ORA analyses of downregulated genes, we also identified enrichments for ribosome and translation for the GO and KEGG databases in physioxic and hypoxic conditions (Fig. 5 A-B). We found that these enrichments are largely specific to FLASH-RT and not associated with the Tumor Growth Delay or All IR overlaps. Similarly, in analyzing the downregulated genes in only the hypoxia condition, we found enrichment for ribosome-and translation-related pathways (Fig. 5 C-D) exclusively in the FLASH only group.

**Fig. 5.**
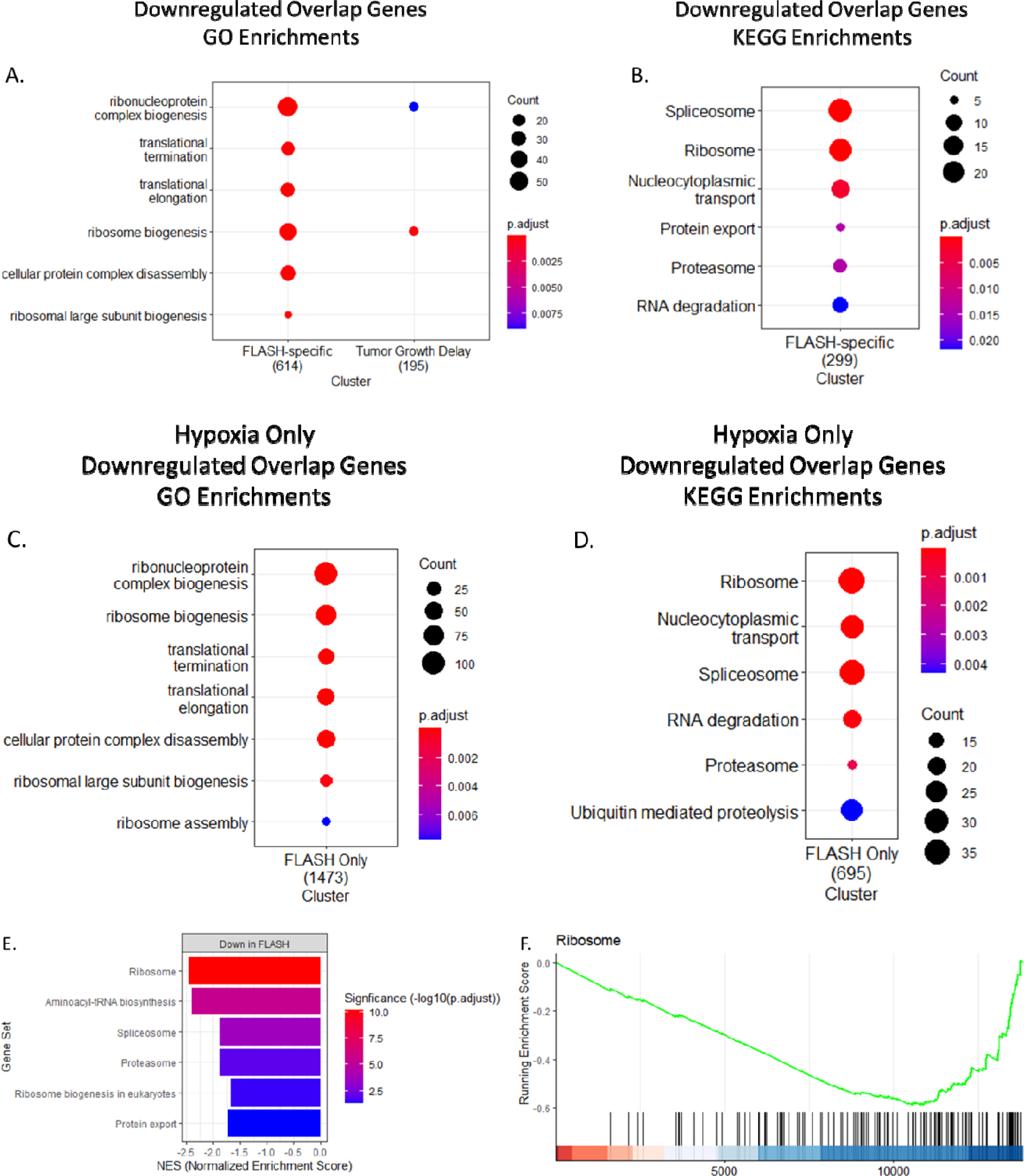
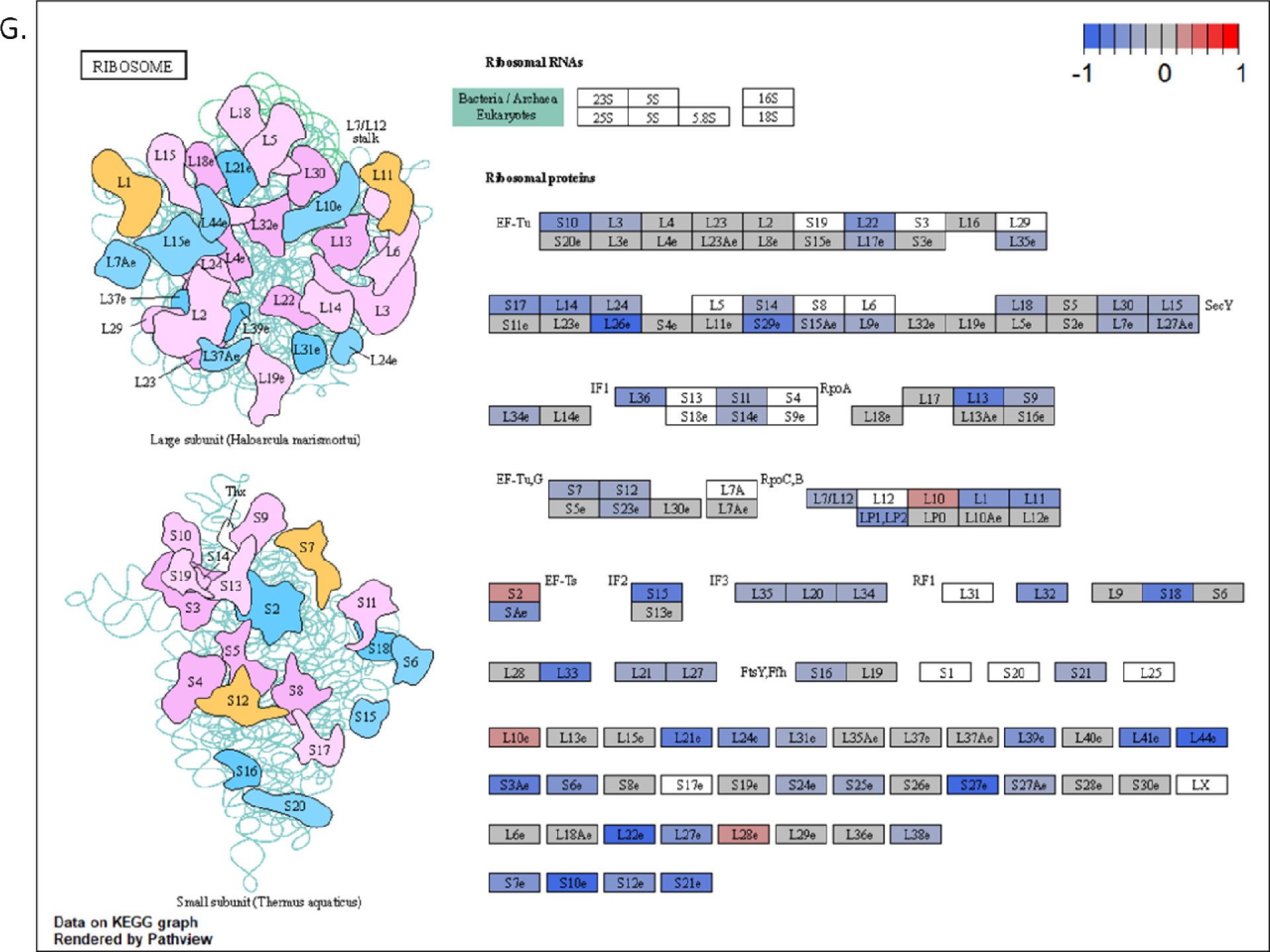
Enrichment of ribosome biogenesis and translation unique to FLASH-RT in both oxygen conditions. Overrepresentation analysis (ORA) for pathways from the Gene Ontology database for differentially expressed gene (DEG) overlap clusters in physioxia (A) and hypoxia (C). ORA for pathways from the Kyoto Encyclopedia of Genes and Genomes (KEGG) database for DEG overlap clusters in physioxia (B) and hypoxia (D). For the hypoxic condition: select ribosome-related pathways (E) enriched in FLASH-RT versus CONV-RT using gene set enrichment analysis (GSEA) for the KEGG database. (F) GSEA plot of the negative (down in FLASH-RT) enrichment for the KEGG ribosome pathway. (G) KEGG ribosome pathway with overlay of the log2 fold changes of FLASH-RT versus CONV-RT (hypoxia) from the DESeq2 output.

Directly comparing the change in gene expression following FLASH-and CONV-RT in hypoxia and determining enrichments using GSEA, we highlighted downregulation of ribosome, mRNA processing, and protein turnover genes, as shown in Fig. 5E and GSEA plot (Fig. 5F). The representation of the ribosome pathway (Fig. 5G) showed widespread downregulation of ribosomal protein transcripts, for both large and small ribosomal subunits.

### Discovery of a FLASH-specific metabolic switch under hypoxic conditions

Using ORA analysis of upregulated genes, we showed enrichment for HIF1 signaling and glycolysis for the Wikipathways and KEGG databases in physioxic and hypoxic conditions (Fig. 6 A-B). These enrichments were specific to the Tumor Growth Delay overlap. The ORA analyses of downregulated genes also showed enrichment for oxidative phosphorylation that was exclusively FLASH-specific. In examining only the hypoxia condition, we found enrichment for HIF1 signaling (up), glycolysis/gluconeogenesis (up), and oxidative phosphorylation (down) only in the FLASH-only overlap (Fig. 6 C-D). Comparing directly the change in gene expression following FLASH-and CONV-RT in hypoxia and determining enrichments using GSEA, we highlighted upregulation of hypoxia and glycolysis and downregulation of oxidative phosphorylation as shown in the Fig. 6E. Examination of the hypoxia and metabolic pathways (Figs. 6, F-H) showed numerous upregulated targets downstream of HIF1A after FLASH hypoxia vs. CONV hypoxia (red, Fig. 6F) – including targets related to anaerobic metabolism such as *GLUT1* (glucose transporter type 1), *HK2* (hexokinase 2), *ALDOA* (aldolase fructose-bisphosphate A), *ENO1* (enolase 1), *PGK1* (phosphoglycerate kinase 1), and *PFK2* (phophofructokinase 2). Specifically, we observed an upregulation of *PDK1* (pyruvate dehydrogenase kinase 1), which, in turn, causes inhibition of the TCA cycle and electron transport chain and stimulation of glycolysis. Therefore, our finding of upregulation of the main glycolytic pathway (red, Fig. 6G) in the FLASH-RT-treated tumors compared to the CONV-RT-treated tumors under hypoxia was consistent with HIF activation. Interestingly, we also found that the section of the pathway responsible for converting pyruvate into acetyl-CoA for the TCA cycle was downregulated (blue), consistent with a lack of aerobic respiration. Additionally, we observed the possibility of the oxaloacetate intermediate being shunted back into the glycolytic/gluconeogenic pathway. Moreover, our analysis revealed that the KEGG oxidative phosphorylation pathway (Fig. 6H) was downregulated in FLASH-versus CONV-RT samples across all five electron transport chain (ETC) complexes, with multiple components downregulated in each of the complex I-V, consistent with significant ETC repression/disruption. To reinforce our ORA and GSEA data, we conducted an unsupervised clustering of all metabolic genes from these two pathways using a heatmap (Fig. 6I), which showed that the genes separated into two clusters - one mostly including OXPHOS genes and another composed mainly of glycolysis/gluconeogenesis genes. We found that the FLASH-RT-treated samples from all oxygen conditions mostly clustered together with the OXPHOS cluster downregulated and the glycolysis cluster upregulated. In contrast, the CONV-RT-treated samples clustered mostly with the non-irradiated ones with higher expression in the OXPHOS cluster and lower expression in the glycolysis one.

**Fig. 6.**
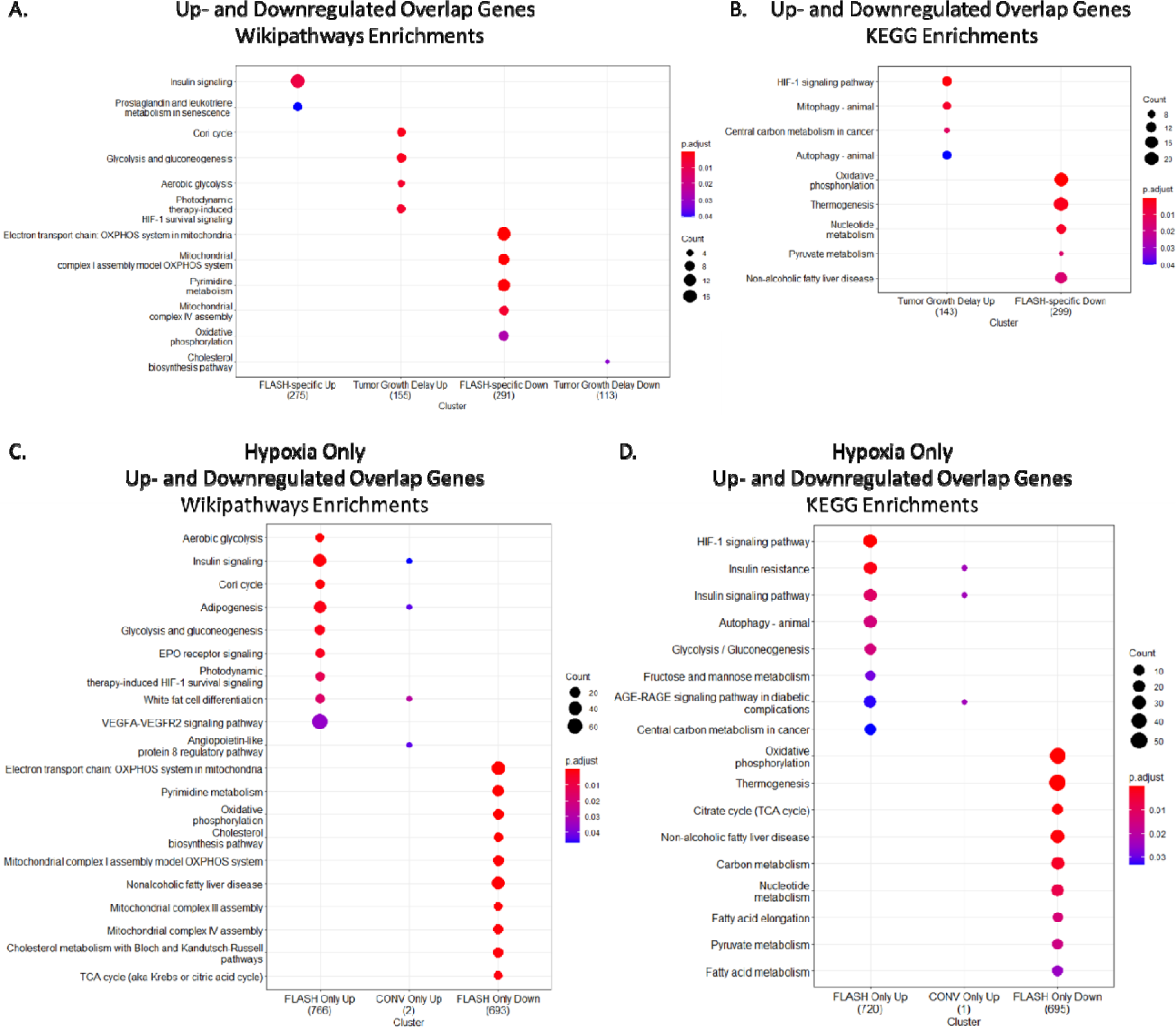
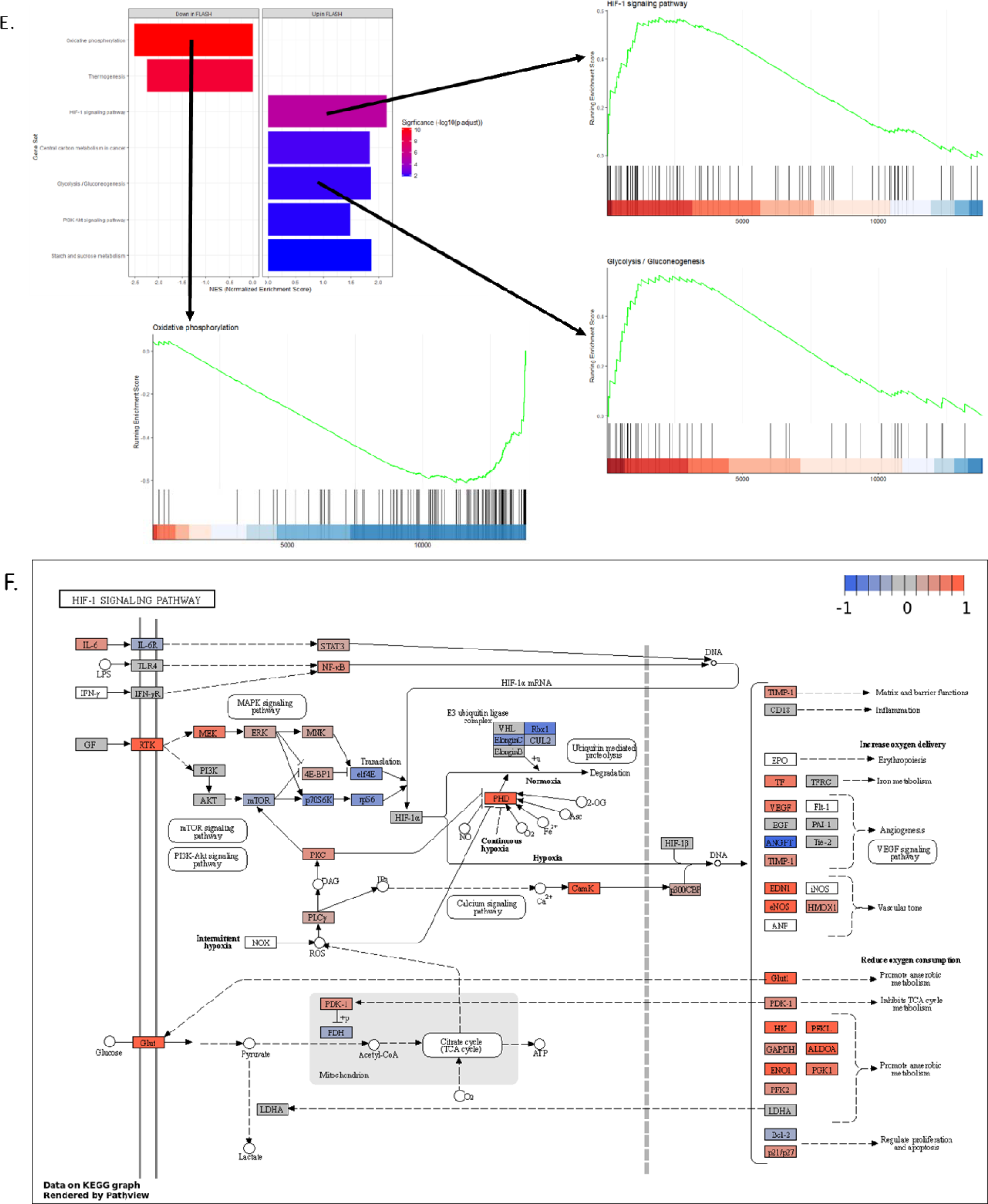
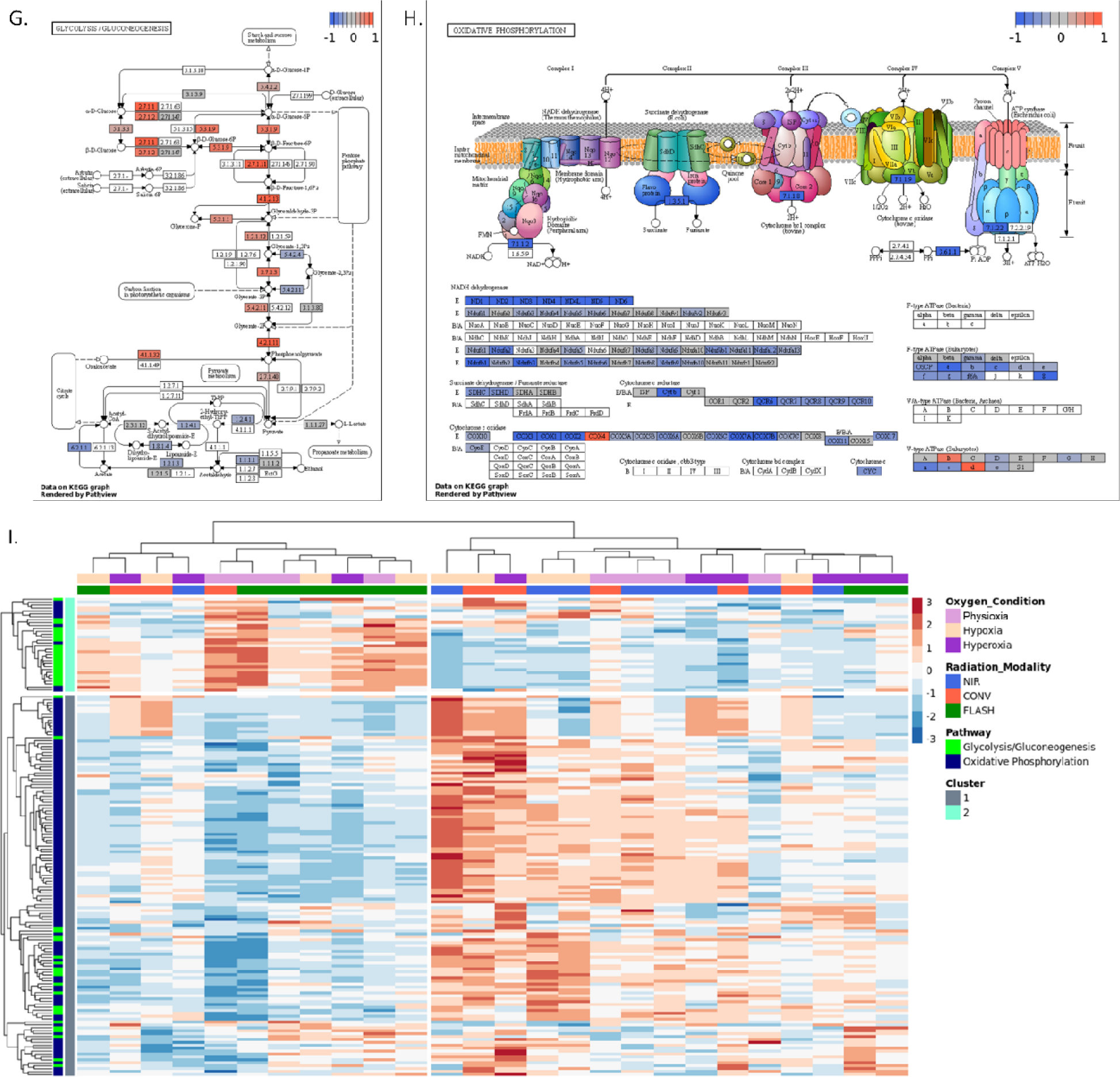
Evidence of FLASH-specific metabolic switch. Overrepresentation analysis (ORA) for pathways from the Wikipathways database for differentially expressed gene (DEG) overlap clusters in physioxia (A) and hypoxia (C). ORA for pathways from the Kyoto Encyclopedia of Genes and Genomes (KEGG) database for DEG overlap clusters in physioxia (B) and hypoxia (D). For the hypoxic condition: select metabolism-related pathways (E) enriched in FLASH-RT versus CONV-RT using gene set enrichment analysis (GSEA) for the KEGG database and GSEA plots of the enrichment for select KEGG pathways. KEGG pathway diagrams of HIF-1 signaling (F), glycolysis/gluconeogenesis (G), and oxidative phosphorylation (H) with overlay of the log2 fold changes of FLASH-RT versus CONV-RT (hypoxia) from the DESeq2 output. (I) Heatmap of exclusively glycolysis/gluconeogenesis and oxidative phosphorylation KEGG pathway genes for all oxygen conditions – columns (samples) and rows (genes, Z-score) clustered hierarchically.

At seven days post-recurrence, PCA (Fig. S6A) and heatmap (Fig. S6B) of the top 500 genes excluding the hyperoxia condition showed a grouping of the non-irradiated samples along with similar profiles (Fig. S6A left and Fig. S6B center-left). The RT treatment groups were not particularly cohesive or homogenous, indicating lots of intra-group sample-to-sample variability. Due to the lack of cohesive group clustering, we could not perform ORA using well-segregated heatmap clusters or lists of genes above a certain log2 fold change cutoff or below an adjusted p-value cutoff. Instead, we looked for signatures of gene sets based on the total set of differentially expressed genes between groups using GSEA and looking for FLASH-specific enrichments (gene sets changed significantly *versus* non-irradiated control in both FLASH physioxia and FLASH hypoxia groups, but not in neither of the CONV-RT groups). Using this analysis, we found that upregulation of metabolic gene sets (mitochondrial genes, mitochondrial translation, and OXPHOS) and the amino acid metabolism gene set were specific to FLASH-RT treated samples at seven days post-recurrence (Fig. S6C). Overall, we suggest that this late time point analysis likely represents the genomic imprint of relapse more so than a CONV-or FLASH-RT response.

These data highlight clear differences in tumor metabolism at early and late time points post-FLASH suggesting that FLASH-induced glycolytic shift in the hypoxia group may be triggering the late relapse and could be targeted to enhance anti-tumor efficacy.

### Neoadjuvant treatment with trametinib

To further evaluate the contribution of glycolysis in tumor relapse following FLASH-RT, we used an FDA-approved MEK 1/2/glycolysis inhibitor called trametinib as an adjuvant treatment to FLASH-RT in hypoxic conditions. We irradiated U-87 MG xenografts with FLASH-RT under hypoxic conditions, combined with a daily oral treatment with 1 mg/kg body weight trametinib starting 24 hours after RT and continuing for two weeks. The adjuvant trametinib treatment delayed the tumor relapse (Fig. 7A), significantly increased the mean doubling time of hypoxic tumors from 32.5 to 40.2 days post-RT (Fig. 7B; P = 0.0054), and improved survival (Fig. 7C; P = 0.0079). Although we observed improved tumor growth delay, our treatment regimen did not result in complete response. These findings suggest that we need to refine the administration schedule and/or investigate the involvement of other resistance mechanisms to enhance the treatment’s efficacy.

**Fig. 7.**
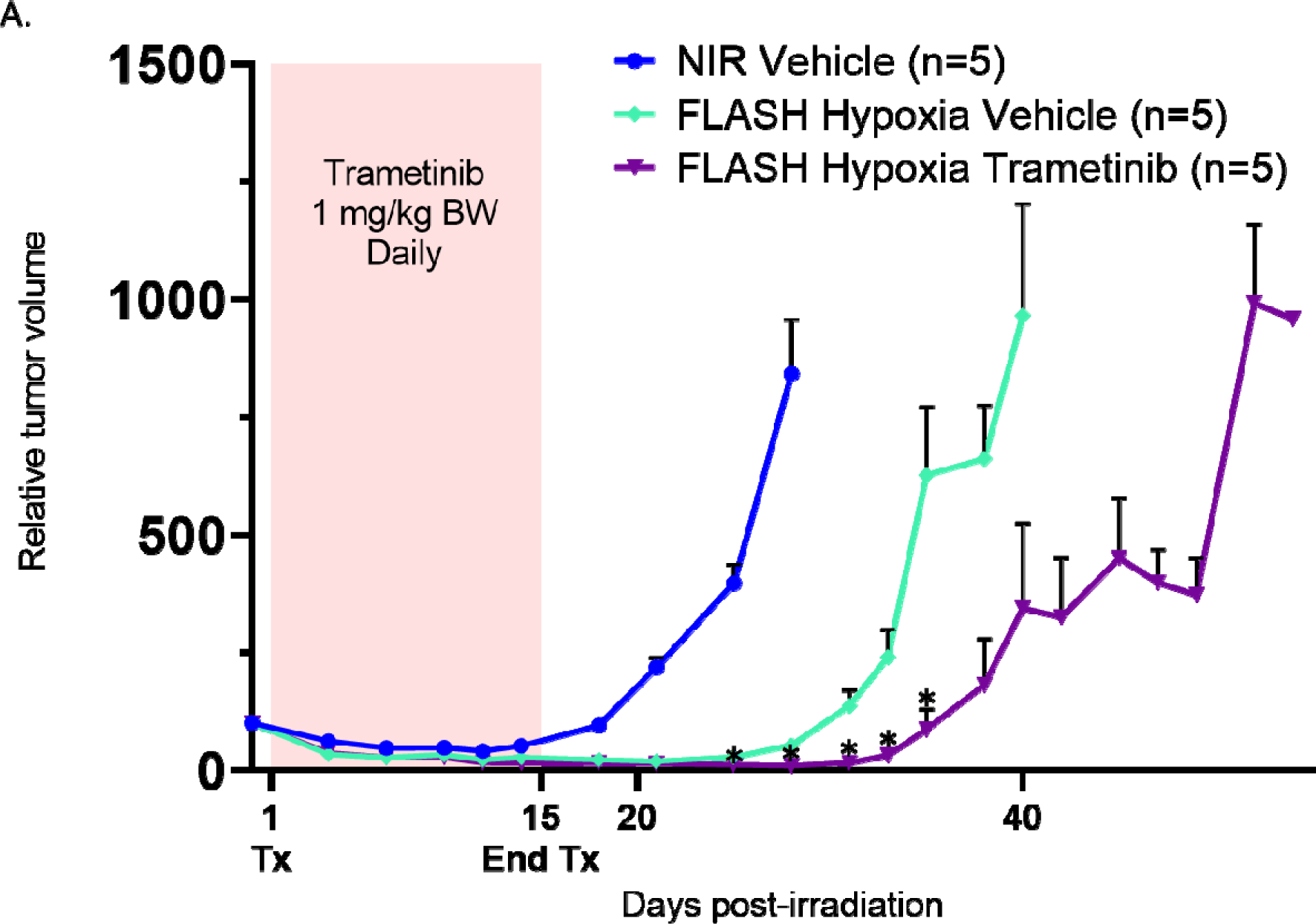
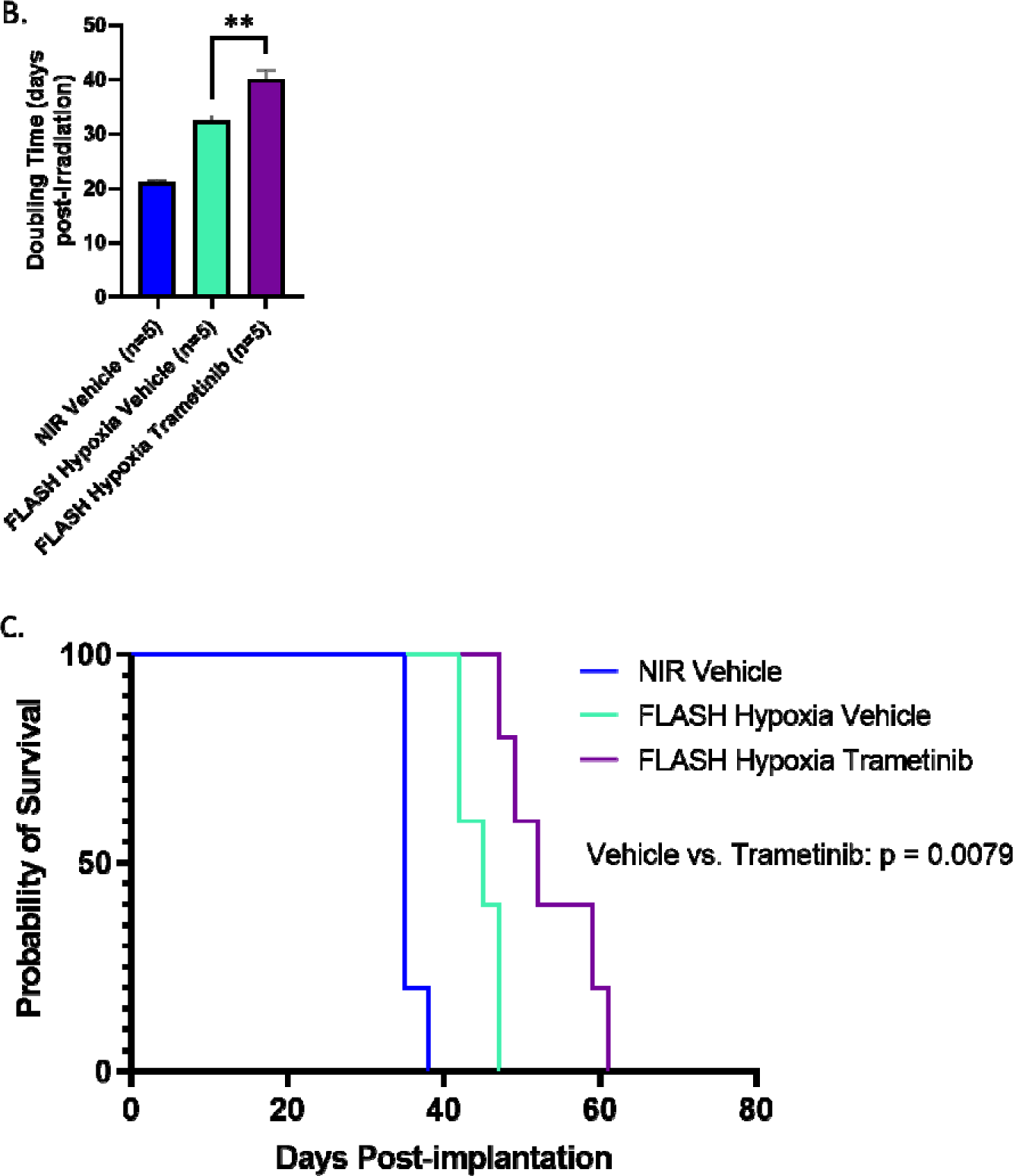
Significant tumor growth delay via daily trametinib treatment for 14 days following FLASH-RT. (A) Relative tumor volume of U-87 MG implanted subcutaneously in the flank of female Nude mice treated with 20 Gy single fraction delivered with FLASH radiation therapy under hypoxic conditions. Mean relative tumor volume + SEM, N = 5 animals per group. P values derived from multiple row-wise Mann-Whitney test against the FLASH-RT hypoxia vehicle group: *P < 0.05. (B) Doubling times for each group. Mean days post-RT + SEM, N=5 animals per group. P values derived from unpaired two-sided t test with Welch’s correction: **P < 0.01. (C) Kaplan–Meier survival curves for animals stratified by radiation modality and tumor oxygenation. P value derived from log rank (Mantel-Cox) test.

## DISCUSSION

As the key finding of this study, we conclusively demonstrated the superior efficacy of FLASH-RT against severely hypoxic tumors compared to CONV-RT. This result was not restricted to a single model as we were able to reproduce and validate it in multiple human and murine tumor models implanted in immunodeficient and immunocompetent mouse strains. Our RNAseq results showed a direct imprint of the response to FLASH vs CONV-RT at acute (24-hour) time-point post-RT, whereas at one week post-recurrence we found the imprint to be relapse-dependent rather than radiation response-dependent. We found evidence of a metabolic switch in both acute and late imprints. Our in-depth analysis of the acute imprint revealed a FLASH-specific profile in hypoxic tumors that involved an arrest of cell cycle, mitosis, and ribosomal biogenesis, and a switch from OXPHOS to glycolysis. Moreover, when we inhibited glycolysis with trametinib, it enhanced FLASH-RT efficacy in the hypoxic condition. Overall, this study highlights the efficacy of FLASH-RT in radiation resistant tumors, mediated by delayed growth and reduced proliferation, and provides a novel if not compelling rationale to use FLASH-RT instead of CONV-RT.

Tumor hypoxia represents one of the major root causes of cancer treatment resistance, and thus hypoxic tumors are extremely difficult to control using current RT, CT, or IT treatments. The relationship between tumor oxygenation and the efficacy of CONV-RT is well-known and has been extensively characterized (4). Our present data obtained with conventional dose rate RT directly reflected this relationship. We found that tumors in hyperoxic conditions experienced a longer growth delay (+ 5 days doubling time vs. physioxic conditions), whereas hypoxic tumors exhibited less favorable control (−8.6 days doubling time vs physioxia). Conversely, FLASH-RT directly challenges this long-standing paradigm with no significant difference in doubling time and growth curves between physioxic and hypoxic tumors. Our results also suggest that the sensitivity of hypoxic tumors to FLASH-RT is not restricted to a single and/or specific model but can be generalized and replicated across various human and murine tumor models. In addition, we observed a similar outcome in immunocompromised and competent hosts, which does not support a specific contribution of the (adaptive) immune system. For rigor, we measured intratumoral oxygen tension directly and non-invasively using the OxyPhor PtG4 probe and OxyLED system. We were able to confirm severe hypoxia (1-2 mmHg O_2_, < 0.5% O_2_) within minutes after clamping (17,23–26), whereas carbogen breathing resulted in an increase in tumor pO_2_ consistent with the literature (27,28). In our study, we showed conclusively that FLASH-RT offers superior control of hypoxic tumors, which are normally resistant to all forms of treatment.

Based on transcriptomic evidence, we suggest that increased tumor control and growth delay using the FLASH modality in hypoxic tumors was due to an inhibition of cell cycle progression caused by specific downregulation of cell cycle genes and proliferation pathways. For the hypoxia condition, we observed downregulation of a number of cell cycle-related genes in both dose-rate groups (Overlap). As a well-characterized response to ionizing radiation and DNA damage (29), we expected it. However, we found an additional cluster of cell cycle-related genes was downregulated in the FLASH only list – with significant independent enrichments for cell cycle and DNA replication. From these data, we surmise that the differential tumor response between FLASH-and CONV-RT under hypoxia is related to a more complete versus partial cell cycle arrest, respectively. Enhanced tumor growth delay observed under FLASH-RT is likely dependent on elevated GADD45. Our observation of increased GADD45 expression in the KEGG Cell Cycle pathway analysis supports this idea. This result is consistent with a putative genomic imprint of FLASH-RT susceptibility shown previously in patient-derived xenograft model of T-cell acute lymphoblastic leukemia (20). The role of GADD45 in controlling activation of S and G2/M checkpoints following genotoxic stress by dissociating and inhibiting the kinase activity of the CDK1/CyclinB1 complex (30) suggests FLASH-RT provides more robust control of proliferation. The increased efficacy of FLASH-RT against the GADD45-overexpressing RKO.AS45.1 tumor xenografts confirmed this result. We infer that the increased *GADD45B* expression found in the U-87 model at 24 hours post-RT under hypoxic conditions indicates cell cycle checkpoints are activated more effectively after FLASH-*vs.* CONV-RT. Interestingly, results also suggest that GADD45 could be used as a prospective biomarker to identify tumor responsiveness to FLASH-RT.

We found that additional gene sets related to translation and ribosomal function were significantly downregulated in FLASH-RT-treated groups in both physioxia and hypoxia. The downregulation of ribosomal machinery is closely associated to cell cycle arrest as previously shown in yeast, in which decreased ribosomal translation was proven to be sufficient to cause cell cycle arrest in the G1 phase (31). Moreover, translation inhibition by cycloheximide caused phosphorylation and activation of CHK1, which impaired G2/M cell cycle progression (32). The translation of cellular mRNAs to proteins by ribosomes can fuel the uncontrolled growth program of a tumor. Additionally, the translation of mRNA into proteins has a high energy cost for cells (33), which is why transient, radiation-induced inhibition of cap-dependent protein synthesis is a conserved cellular stress response (34). Our results suggest that FLASH-RT enhances the level of ribosomal stress in tumors, thus decreasing translation, which can induce further activation of cell cycle checkpoints, thereby overriding the protective effects of hypoxia.

Next, we showed that the tumor growth delay outcome under severe hypoxia was associated with significant metabolic alterations 24 hours post-FLASH-RT, and more specifically with an increased glycolysis and decreased OXPHOS gene expression. These changes are consistent with recently described alterations in cancer cell metabolism following *in vitro* exposure to ionizing radiation (35). However, we did not detect such changes upon comparison of the CONV-RT hypoxia to the group to the NIR hypoxia group, indicating that FLASH-RT modulates the expression of these metabolic gene sets under severe hypoxia, while CONV-RT does not. We presented overwhelming evidence of these metabolic changes using ORA, GSEA, pathway analysis, and heatmap clustering of metabolic genes. Furthermore, we found that OXPHOS/mitochondrial gene sets were significantly upregulated in FLASH-RT-treated tumors (Fig. S6) under both hypoxia and physioxia conditions seven days post-recurrence. These results correlate with studies showing a higher treatment resistance in OXYPHOS-dependent, dormant, and slow-cycling tumor stem cells (36–38). However, the fact that we observed greater enrichment of these gene sets in FLASH-versus CONV-RT treated tumors suggests that FLASH-RT elicits a stronger metabolic selection. In summary, we discovered a FLASH-RT-specific metabolic switch from OXPHOS to glycolysis under hypoxia that was reversed at latter times of tumor relapse.

The metabolic shift prompted us to test a glycolysis inhibitor, trametinib. Trametinib is a FDA-approved MEK1/2 inhibitor, that can be administered orally and is well tolerated (39). Results from a recent clinical trial provided evidence that that the drug could readily cross the blood-brain barrier in humans (40), which is essential for GBM treatment. U-87 MG cells were previously shown to be sensitive to trametinib (41) and this drug was found to inhibit growth (*in vitro* and *in vivo*) and aerobic glycolysis (*in vitro*) in other glioma models (42). These features clearly boost the clinical relevance of our study, where we used trametinib as a short-term adjuvant treatment and showed its ability to enhance the anti-tumor efficacy of FLASH-RT in the U-87 MG xenograft model under hypoxia. This key evidence suggests that the combination of cell cycle arrest and metabolic intervention are key factors that are involved and can be exploited to optimize FLASH efficacy.

So far, the superiority of FLASH-over CONV-RT has largely been attributed to its capability to spare normal tissue from radiation-induced toxicities. In the present study, we brought the first evidence of an important additional benefit of FLASH-RT, namely, efficacy against radioresistant hypoxic tumors compared to conventional dose rate RT. We complemented these findings with the identification of a FLASH-induced inhibition of proliferation and translation. Notably we also uncovered a druggable metabolic switch that we successfully reversed using the FDA-approved compound trametinib to enhance the efficacy of FLASH-RT. These mechanistic findings are fundamental and support the ultimate translation of FLASH-RT into the clinic.

## Acknowledgments

The authors thank Prs J Bourhis and F Bochud for their support, Dr S Vinogradov for expert technical advice, Dr T Boehlen for help in formatting figures, J Ollivier for technical help, K Sprengers for help in radiation experiments. We also thank the Lausanne Genomic Technologies Facility and the animal facility at Epalinges.

## References

1. Vasan N, Baselga J, Hyman DM. A view on drug resistance in cancer. Nature. 2019;575:299–309.

2. Vaupel P, Höckel M, Mayer A. Detection and Characterization of Tumor Hypoxia Using pO2 Histography. Antioxidants & Redox Signaling. Mary Ann Liebert, Inc., publishers; 2007;9:1221–36.

3. Zhu W, Dong Z, Fu T, Liu J, Chen Q, Li Y, et al. Modulation of Hypoxia in Solid Tumor Microenvironment with MnO2 Nanoparticles to Enhance Photodynamic Therapy. Advanced Functional Materials. 2016;26:5490–8.

4. Moulder JE, Rockwell S. Hypoxic fractions of solid tumors: Experimental techniques, methods of analysis, and a survey of existing data. International Journal of Radiation Oncology*Biology*Physics. 1984;10:695– 712.

5. Zeng W, Liu P, Pan W, Singh SR, Wei Y. Hypoxia and hypoxia inducible factors in tumor metabolism. Cancer Letters. 2015;356:263–7.

6. Graham K, Unger E. Overcoming tumor hypoxia as a barrier to radiotherapy, chemotherapy and immunotherapy in cancer treatment. IJN. 2018;Volume 13:6049–58.

7. Favaudon V, Caplier L, Monceau V, Pouzoulet F, Sayarath M, Fouillade C, et al. Ultrahigh dose-rate FLASH irradiation increases the differential response between normal and tumor tissue in mice. Sci Transl Med. 2014;6:245ra93.

8. Montay-Gruel P, Petersson K, Jaccard M, Boivin G, Germond JF, Petit B, et al. Irradiation in a flash: Unique sparing of memory in mice after whole brain irradiation with dose rates above 100Gy/s. Radiother Oncol. 2017;124:365–9.

9. Montay-Gruel P, Bouchet A, Jaccard M, Patin D, Serduc R, Aim W, et al. X-rays can trigger the FLASH effect: Ultra-high dose-rate synchrotron light source prevents normal brain injury after whole brain irradiation in mice. Radiother Oncol. 2018;129:582–8.

10. Vozenin MC, De Fornel P, Petersson K, Favaudon V, Jaccard M, Germond JF, et al. The Advantage of FLASH Radiotherapy Confirmed in Mini-pig and Cat-cancer Patients. Clin Cancer Res. 2019;25:35–42.

11. Montay-Gruel P, Acharya MM, Petersson K, Alikhani L, Yakkala C, Allen BD, et al. Long-term neurocognitive benefits of FLASH radiotherapy driven by reduced reactive oxygen species. Proc Natl Acad Sci U S A. 2019;116:10943–51.

12. Bourhis J, Sozzi WJ, Jorge PG, Gaide O, Bailat C, Duclos F, et al. Treatment of a first patient with FLASH-radiotherapy. Radiotherapy and Oncology. 2019;139:18–22.

13. Alaghband Y, Cheeks SN, Allen BD, Montay-Gruel P, Doan NL, Petit B, et al. Neuroprotection of Radiosensitive Juvenile Mice by Ultra-High Dose Rate FLASH Irradiation. Cancers (Basel). 2020;12.

14. Vozenin M-C, Bourhis J, Durante M. Towards clinical translation of FLASH radiotherapy. Nat Rev Clin Oncol. Nature Publishing Group; 2022;19:791–803.

15. Vozenin M-C, Hendry JH, Limoli CL. Biological Benefits of Ultra-high Dose Rate FLASH Radiotherapy: Sleeping Beauty Awoken. Clinical Oncology. 2019;31:407–15.

16. Wilson JD, Hammond EM, Higgins GS, Petersson K. Ultra-High Dose Rate (FLASH) Radiotherapy: Silver Bullet or Fool’s Gold? Frontiers in Oncology [Internet]. 2020 [cited 2022 Apr 12];9. Available from: https://www.frontiersin.org/article/10.3389/fonc.2019.01563

17. Cao X, Zhang R, Esipova TV, Allu SR, Ashraf R, Rahman M, et al. Quantification of Oxygen Depletion During FLASH Irradiation In Vitro and In Vivo. International Journal of Radiation Oncology*Biology*Physics. 2021;111:240–8.

18. Pratx G, Kapp DS. Ultra-High-Dose-Rate FLASH Irradiation May Spare Hypoxic Stem Cell Niches in Normal Tissues. International Journal of Radiation Oncology*Biology*Physics. 2019;105:190–2.

19. Fouillade C, Curras-Alonso S, Giuranno L, Quelennec E, Heinrich S, Bonnet-Boissinot S, et al. FLASH Irradiation Spares Lung Progenitor Cells and Limits the Incidence of Radio-induced Senescence. Clinical Cancer Research. 2020;26:1497–506.

20. Chabi S, To THV, Leavitt R, Poglio S, Jorge PG, Jaccard M, et al. Ultra-high-dose-rate FLASH and Conventional-Dose-Rate Irradiation Differentially Affect Human Acute Lymphoblastic Leukemia and Normal Hematopoiesis. International Journal of Radiation Oncology*Biology*Physics. 2021;109:819–29.

21. Ruan J-L, Lee C, Wouters S, Tullis IDC, Verslegers M, Mysara M, et al. Irradiation at Ultra-High (FLASH) Dose Rates Reduces Acute Normal Tissue Toxicity in the Mouse Gastrointestinal System. International Journal of Radiation Oncology*Biology*Physics. 2021;111:1250–61.

22. Montay-Gruel P, Acharya MM, Gonçalves Jorge P, Petit B, Petridis IG, Fuchs P, et al. Hypofractionated FLASH-RT as an Effective Treatment against Glioblastoma that Reduces Neurocognitive Side Effects in Mice. Clinical Cancer Research. 2021;27:775–84.

23. Dunphy I, Vinogradov SA, Wilson DF. Oxyphor R2 and G2: phosphors for measuring oxygen by oxygen-dependent quenching of phosphorescence. Analytical Biochemistry. 2002;310:191–8.

24. Esipova TV, Karagodov A, Miller J, Wilson DF, Busch TM, Vinogradov SA. Two New “Protected” Oxyphors for Biological Oximetry: Properties and Application in Tumor Imaging. Anal Chem. American Chemical Society; 2011;83:8756–65.

25. Esipova TV, Barrett MJP, Erlebach E, Masunov AE, Weber B, Vinogradov SA. Oxyphor 2P: A High-Performance Probe for Deep-Tissue Longitudinal Oxygen Imaging. Cell Metabolism. 2019;29:736–744.e7.

26. El Khatib M, Van Slyke AL, Velalopoulou A, Kim MM, Shoniyozov K, Allu SR, et al. Ultrafast Tracking of Oxygen Dynamics During Proton FLASH. International Journal of Radiation Oncology*Biology*Physics. 2022;113:624–34.

27. Martin L, Lartigau E, Weeger P, Lambin P, Le Ridant AM, Lusinchi A, et al. Changes in the oxygenation of head and neck tumors during carbogen breathing. Radiotherapy and Oncology. 1993;27:123–30.

28. Falk SJ, Ward R, Bleehen NM. The influence of carbogen breathing on tumour tissue oxygenation in man evaluated by computerised p02 histography. Br J Cancer. Nature Publishing Group; 1992;66:919–24.

29. Visconti R, Della Monica R, Grieco D. Cell cycle checkpoint in cancer: a therapeutically targetable double-edged sword. Journal of Experimental & Clinical Cancer Research. 2016;35:153.

30. Vairapandi M, Balliet AG, Hoffman B, Liebermann DA. GADD45b and GADD45g are cdc2/cyclinB1 kinase inhibitors with a role in S and G2/M cell cycle checkpoints induced by genotoxic stress. Journal of Cellular Physiology. 2002;192:327–38.

31. Shamsuzzaman M, Bommakanti A, Zapinsky A, Rahman N, Pascual C, Lindahl L. Analysis of cell cycle parameters during the transition from unhindered growth to ribosomal and translational stress conditions. PLOS ONE. Public Library of Science; 2017;12:e0186494.

32. Thomas SE, Malzer E, Ordóñez A, Dalton LE, Wout EFA van ′t, Liniker E, et al. p53 and Translation Attenuation Regulate Distinct Cell Cycle Checkpoints during Endoplasmic Reticulum (ER) Stress *. Journal of Biological Chemistry. Elsevier; 2013;288:7606–17.

33. Li G-W, Burkhardt D, Gross C, Weissman JS. Quantifying Absolute Protein Synthesis Rates Reveals Principles Underlying Allocation of Cellular Resources. Cell. 2014;157:624–35.

34. Braunstein S, Badura ML, Xi Q, Formenti SC, Schneider RJ. Regulation of Protein Synthesis by Ionizing Radiation. Molecular and Cellular Biology. American Society for Microbiology; 2009;29:5645–56.

35. Krysztofiak A, Szymonowicz K, Hlouschek J, Xiang K, Waterkamp C, Larafa S, et al. Metabolism of cancer cells commonly responds to irradiation by a transient early mitochondrial shutdown. iScience. 2021;24:103366.

36. Viale A, Pettazzoni P, Lyssiotis CA, Ying H, Sánchez N, Marchesini M, et al. Oncogene ablation-resistant pancreatic cancer cells depend on mitochondrial function. Nature. Nature Publishing Group; 2014;514:628– 32.

37. Lagadinou ED, Sach A, Callahan K, Rossi RM, Neering SJ, Minhajuddin M, et al. BCL-2 Inhibition Targets Oxidative Phosphorylation and Selectively Eradicates Quiescent Human Leukemia Stem Cells. Cell Stem Cell. 2013;12:329–41.

38. Viale A, Corti D, Draetta GF. Tumors and Mitochondrial Respiration: A Neglected Connection. Cancer Research. 2015;75:3687–91.

39. Wright CJ, McCormack PL. Trametinib: first global approval. Drugs. 2013;73:1245–54.

40. Wen PY, Stein A, Bent M van den, Greve JD, Wick A, Vos FYFL de, et al. Dabrafenib plus trametinib in patients with BRAFV600E-mutant low-grade and high-grade glioma (ROAR): a multicentre, open-label, single-arm, phase 2, basket trial. The Lancet Oncology. Elsevier; 2022;23:53–64.

41. Kanemaru Y, Natsumeda M, Okada M, Saito R, Kobayashi D, Eda T, et al. Dramatic response of BRAF V600E-mutant epithelioid glioblastoma to combination therapy with BRAF and MEK inhibitor: establishment and xenograft of a cell line to predict clinical efficacy. Acta Neuropathologica Communications. 2019;7:119.

42. Gao M, Yang J, Gong H, Lin Y, Liu J. Trametinib Inhibits the Growth and Aerobic Glycolysis of Glioma Cells by Targeting the PKM2/c-Myc Axis. Frontiers in Pharmacology [Internet]. 2021 [cited 2022 Jul 19];12. Available from: https://www.frontiersin.org/articles/10.3389/fphar.2021.760055

43. Gene Expression Omnibus: NCBI gene expression and hybridization array data repository | Nucleic Acids Research | Oxford Academic [Internet]. [cited 2023 Jan 24]. Available from: https://academic.oup.com/nar/article/30/1/207/1332640?login=true

