## Supplementary material for "Hypoxic tumors are sensitive to FLASH radiotherapy": supp file

SUPPLEMENTARY MATERIALS

**Supplementary Methods**

Irradiation device and beam parameters

Irradiations were performed using a prototype 6 MeV electron beam linear accelerator of type Oriatron 6e (eRT6; PMB Alcen), available at Lausanne University Hospital and described previously (1). Dosimetry has been extensively described and published to ensure reproducible and reliable biological studies. All FLASH irradiations were performed at a mean dose rate greater than or equal to 100 Gy/s and at an intra-pulse dose rate above 5.6x10^6^ Gy/s. The beam parameters used throughout this study are included in Table 1. The irradiation settings corresponding to the prescription dose for mouse irradiations were determined by surface dose measurements on a 30 × 30 cm^2^-solid water slab positioned behind a 1.7 cm-diameter aperture of a graphite applicator (13.0 × 13.0 × 2.5 cm^3^), as previously described.

**Table 1. Oriatron eRT6 beam parameters in FLASH and CONV modes**

| Mode | Prescribed dose (Gy) | Frequency (Hz) | Grid tension (V) | Number of pulses | Treatment time (s) | Pulse Width (μs) | Mean Dose Rate (Gy/s) | Instantaneous Dose Rate (Gy/s) |
| --- | --- | --- | --- | --- | --- | --- | --- | --- |
| FLASH | 20 | 100 | 300 | 2-20 | 0.01-0.2 | 1.8 | 100 - 1993 | 5.56 x 10^5^ – 5.56 x 10^6^ |
| CONV | 20 | 10 | 100 | 1942 | 194.1 | 1 | 0.1 | 1.03 x 10^4^ |

Cell culture

U-87 MG (HTB-14, ATCC, Germany) human glioblastoma, murine H454 glioblastoma, SV2 lung adenocarcinoma, and human RKO (CRL-2577, ATCC, Germany) and RKO.AS45.1 (CRL-2579, ATCC, Germany) colon carcinoma cell lines were cultured in complete medium containing DMEM (Dulbecco’s Modified Eagle Medium) + GlutaMAX (4.5 g/L D-Glucose, Pyruvate; 31966-021, Thermo Fisher Scientific) and supplemented with 10% FBS (F7524, Sigma-Aldrich).

mEERL95 murine head and neck squamous cell carcinoma cells were cultured in MDMEM/Medium F-12 + GlutaMAX (31331028, Thermo Fisher Scientific), 5% FBS (F7524, Sigma), penicillin/streptomycin (15140122 Thermo Fisher Scientific) and human keratinocyte growth supplement (HKGS, S-001-5, Thermo Fisher Scientific).

Cell lines were maintained in an incubator at 37°C, 5% CO2 and routinely tested to dismiss Mycoplasma infection.

Tumor Volume and Relative Tumor Volume Calculations

The tumor volume was measured 3x / week using a digital caliper and calculated using the formula for an oblate ellipsoid: $\frac{{width}^{2}\times length}{2}$. Relative tumor volumes were calculated by percent of volume at the time of irradiation ($\frac{Current Volume}{Volume at T_{0}} \times100\%)$.

Anesthesia of animals

All irradiations of subcutaneous tumors were performed under isoflurane anesthesia. Induction was performed in an induction box with 3-4% isoflurane, and anesthesia was maintained using a mask with 1-2% isoflurane concentration. Animals were placed on a heating mat to maintain their body temperature at 37°C. Vitamin A in ophthalmic cream (Vitapos®) was applied to the eyes of the animal to prevent drying.

Validation of the different oxygen conditions with Pimonidazole

To validate the different oxygenation conditions of the tumors, mice from each oxic condition group (n=4 / group) were injected intravenously with 60 mg/kg body weight Pimonidazole (HP1-100Kit, Hypoxyprobe), 90 min before tumor sampling. For physioxic conditions, injection was followed by 90 min normal air breathing. For hyperoxic conditions, injection was followed by 90 min carbogen breathing. For hypoxic conditions, injection was followed by 15 min air breathing followed by 7 min tumor clamping and finally 68 min air breathing.

Tumors were then collected, fixed in FineFix (84-1717-00, Biosystems), embedded in paraffin and finally cut into 4 µm sections. Tumor hypoxia was validated on tumor sections using mouse anti-pimonidazole monoclonal antibody (1:50; HP1-100Kit, Hypoxyprobe) incubated 1h at room temperature. The sections were then incubated for 1h with a donkey anti-mouse AF488 secondary antibody (1:250; A21202, Life Technologies). Image acquisition was performed using an upright Zeiss Axiovision microscope.

RNA extraction and High-Throughput Sequencing

For the RNAseq study, mice (n=3-6 for each group) were irradiated in the same conditions and with the same dose as described above. Tumors were sampled 24 hours after radiotherapy or 7 days post-recurrence and were disrupted in cold RLT lysis buffer with automated homogenizer (TissueLyser II, Qiagen) and total RNA was extracted using the RNEasy mini kit (Qiagen) according to manufacturer instructions. RNA quality (RQN > 6.8) was assessed on a Fragment Analyzer (Agilent Technologies). RNAseq libraries were prepared from 500 ng of total RNA with the Illumina TruSeq Stranded mRNA reagents (Illumina) using a unique dual indexing strategy, and following the official protocol automated on a Sciclone liquid handling robot (PerkinElmer). Libraries were quantified by a fluorimetric method (QubIT, Life Technologies) and their quality assessed on a Fragment Analyzer (Agilent Technologies).

Cluster generation was performed with 2 nM of an equimolar pool from the resulting libraries using the Illumina HiSeq 3000/4000 SR Cluster Kit reagents and sequenced on the Illumina HiSeq 4000 using HiSeq 3000/4000 SBS Kit reagents for 150 cycles (single end).

Bioinformatic Analysis

Raw FASTQ files were uploaded to the European Galaxy server (www.usegalaxy.eu) for further manipulation and processing. Read quality was assessed using FastQC. Read trimming was not necessary (2) thus alignment was immediately performed. The RNA STAR aligner (version 2.7.8a, (3)) was used to align the reads to the human genome (hg38, for the human tumor cells) and the mouse genome (mm10, for the mouse vasculature within the tumor sample), with NM-tag turned on for subsequent XenofilteR analysis. Binary alignment (BAM) files from all sequencing lanes for each sample were merged at this point (within alignments to the same species) using the Samtools merge tool (version 1.13, (4)). These merged and paired alignments for each sample were used as input for XenofilteR (version 1.6, (5)) for filtering of mouse reads from the human alignment, which was implemented in R (version 4.1.0, (6)) using the RStudio environment (version 1.4.1717, (7)). These filtered reads were then counted for annotated genes using featureCounts (version 2.0.1, (8)) in Galaxy. Raw counts tables were imported back into RStudio for differential gene expression and pathway analyses using the DESeq2 (version 1.34.0, (9)) and clusterProfiler (version 4.2.0, (10)) packages. Plots were generated using the ggplots2 (version 3.3.5, (11)), pheatmap (version 1.0.12, (12)), and Pathview (version 1.32.0, (13)) packages.

Expanded Statistical Analyses

Growth curves were not normally distributed, thus P values comparing tumor growth delay curves were derived from multiple row-wise Mann-Whitney U tests. Tumor doubling times were calculated by natural-log-transformation of relative tumor volumes, followed by line fitting (only log-linear range), followed by interpolation/extrapolation of the point where the fitted line crosses the doubling threshold. Standard deviations were not consistent among groups; therefore, P values for group comparisons of doubling time were derived from unpaired t tests with Welch’s correction following Brown-Forsythe and Welch one-way ANOVA tests. P values comparing survival curves were derived from the log rank (Mantel-Cox) test. For RNAseq analysis, all adjusted p-values had false discovery rate (FDR) correction applied using the Benjamini-Hochberg procedure (14).

**Supplementary Figures**

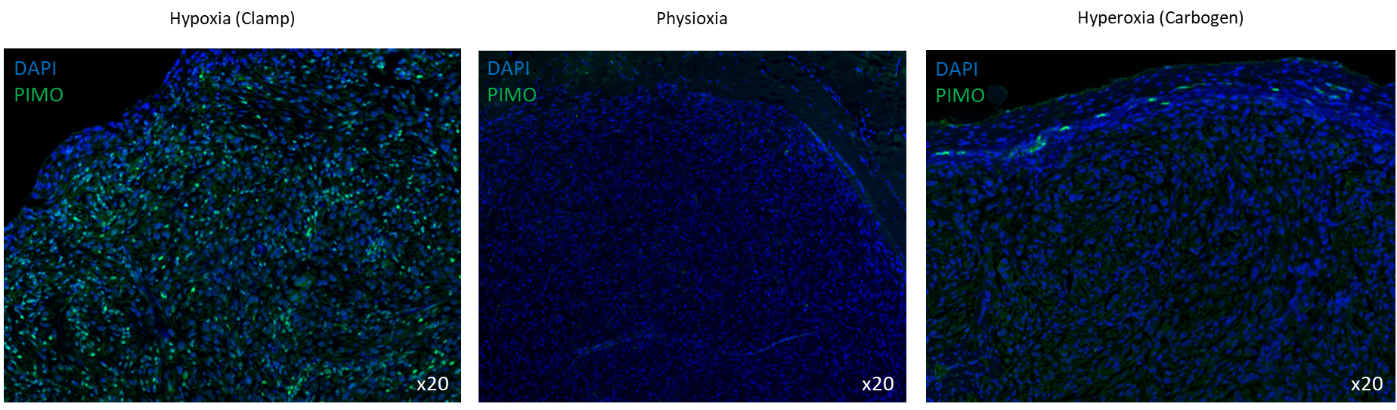

**Fig. S1.** Pimonidazole immunostaining on tumor sections after manipulating oxygenation conditions: physioxia, vascular clamp, or carbogen breathing – each without radiotherapy. Green, Pimonidazole (PIMO); Blue, DAPI.

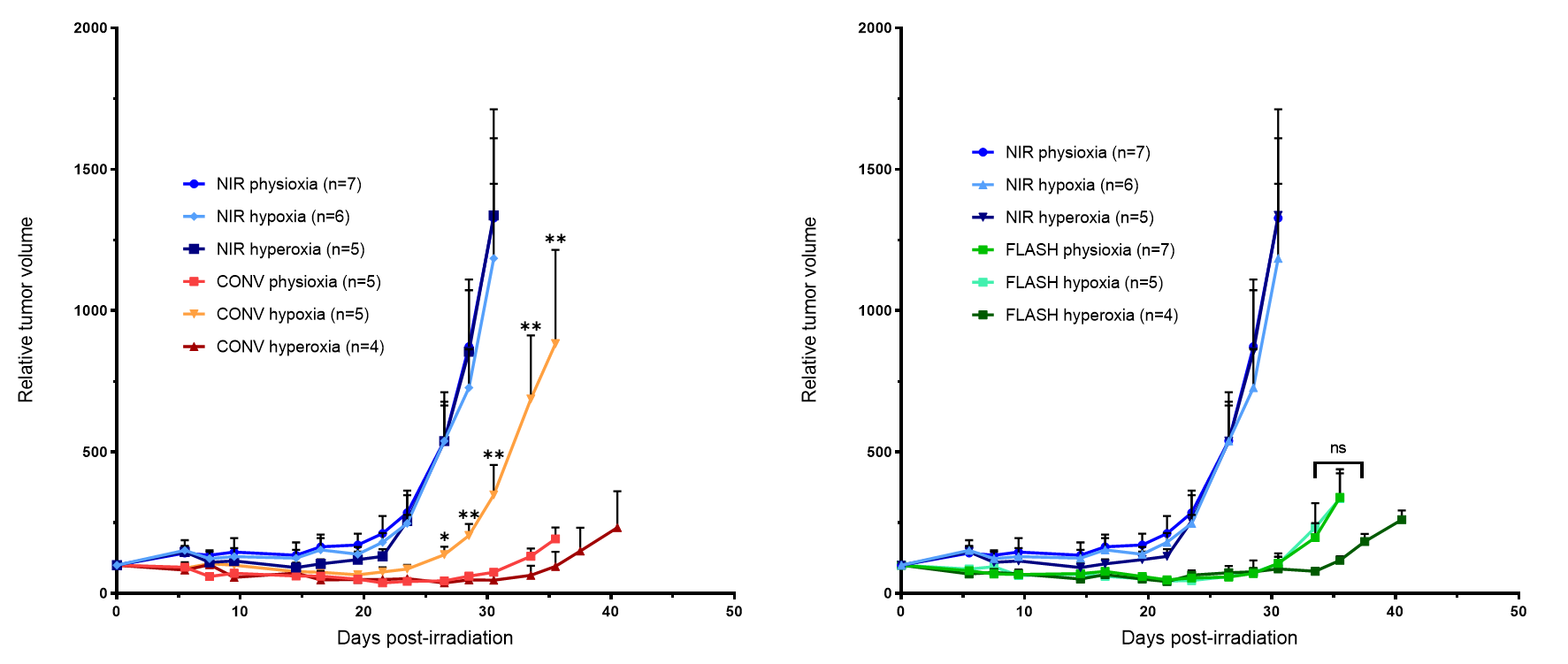

**Fig. S2.** Relative tumor volume of U-87 MG human GBM cells implanted in subcutaneous of female Nude mice treated with 20 Gy single fraction delivered with CONV (left) or FLASH (right) radiation therapy with different oxygenation conditions (physioxia – hypoxia – hyperoxia). Animal cohort sampled for the late time point RNAseq analysis. Mean relative tumor volume + SEM, N = 4-7 animals per group. P values were derived from multiple row-wise Mann-Whitney tests against the FLASH physioxia group: *P < 0.05; **P < 0.01; ns, not significant.

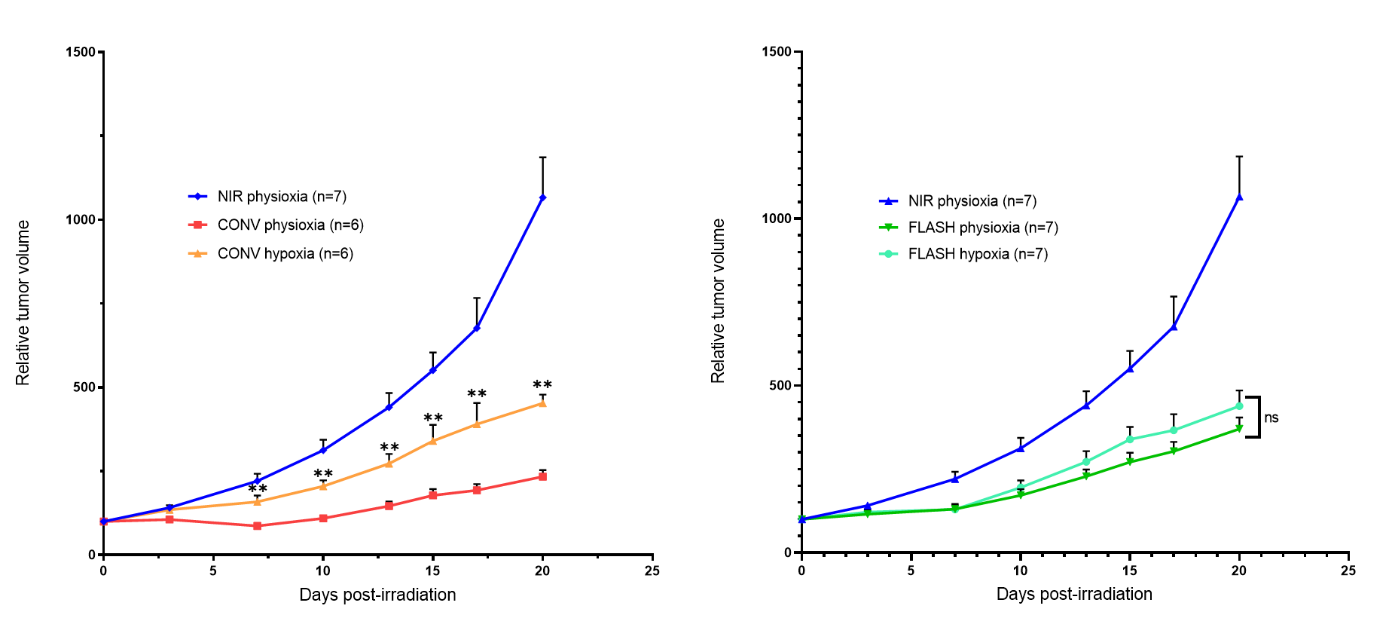

**Fig. S3.** Relative tumor volume of H454 mouse GBM cells implanted in subcutaneous of female Nude mice treated with 20 Gy single fraction delivered with CONV (left) or FLASH (right) radiation therapy with different oxygenation conditions (physioxia – hypoxia – hyperoxia). Mean relative tumor volume + SEM, N = 6-7 animals per group. P values were derived from multiple row-wise Mann-Whitney tests against the FLASH physioxia group: **P < 0.01; ns, not significant.

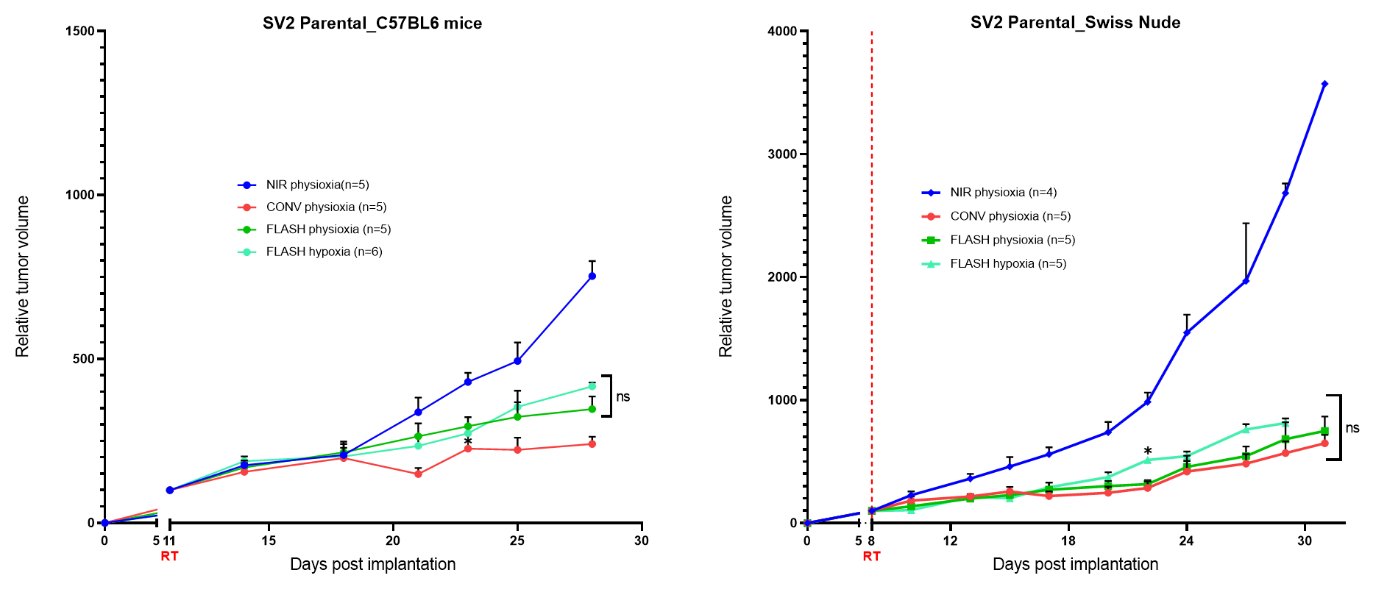

**Fig. S4.** Relative tumor volume of SV2 mouse lung adenocarcinoma cells implanted in subcutaneous of female C57BL/6J (left) or Swiss nude (right) mice treated with 20 Gy single fraction delivered with CONV or FLASH radiation therapy with different oxygenation conditions (physioxia – hypoxia). Mean relative tumor volume + SEM, N = 4-6 animals per group. P values were derived from multiple row-wise Mann-Whitney tests against the FLASH physioxia group: *P < 0.05; ns, not significant.

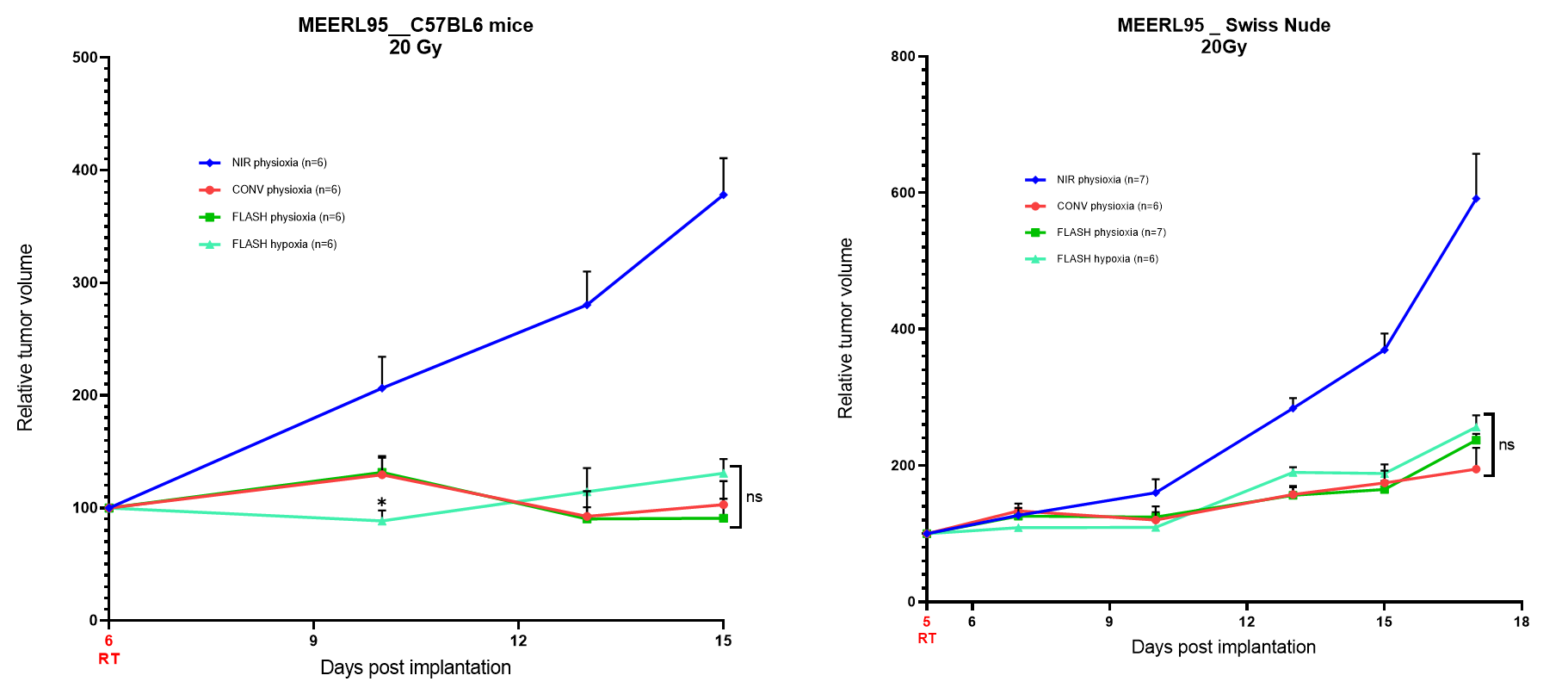

**Fig. S5.** Relative tumor volume of MEERL95 mouse head and neck carcinoma cells implanted in subcutaneous of female C57BL/6JRJ (left) or Swiss nude (right) mice treated with 20 Gy single fraction delivered with CONV or FLASH radiation therapy with different oxygenation conditions (physioxia – hypoxia). Mean relative tumor volume + SEM, N = 6-7 animals per group. P values were derived from multiple row-wise Mann-Whitney tests against the FLASH physioxia group: *P < 0.05; ns, not significant.

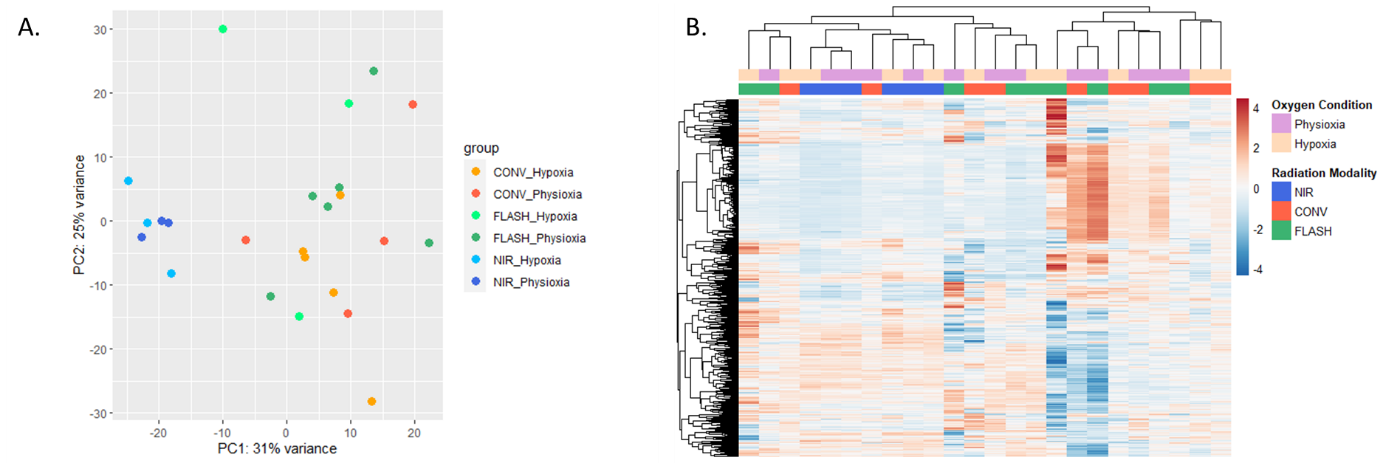

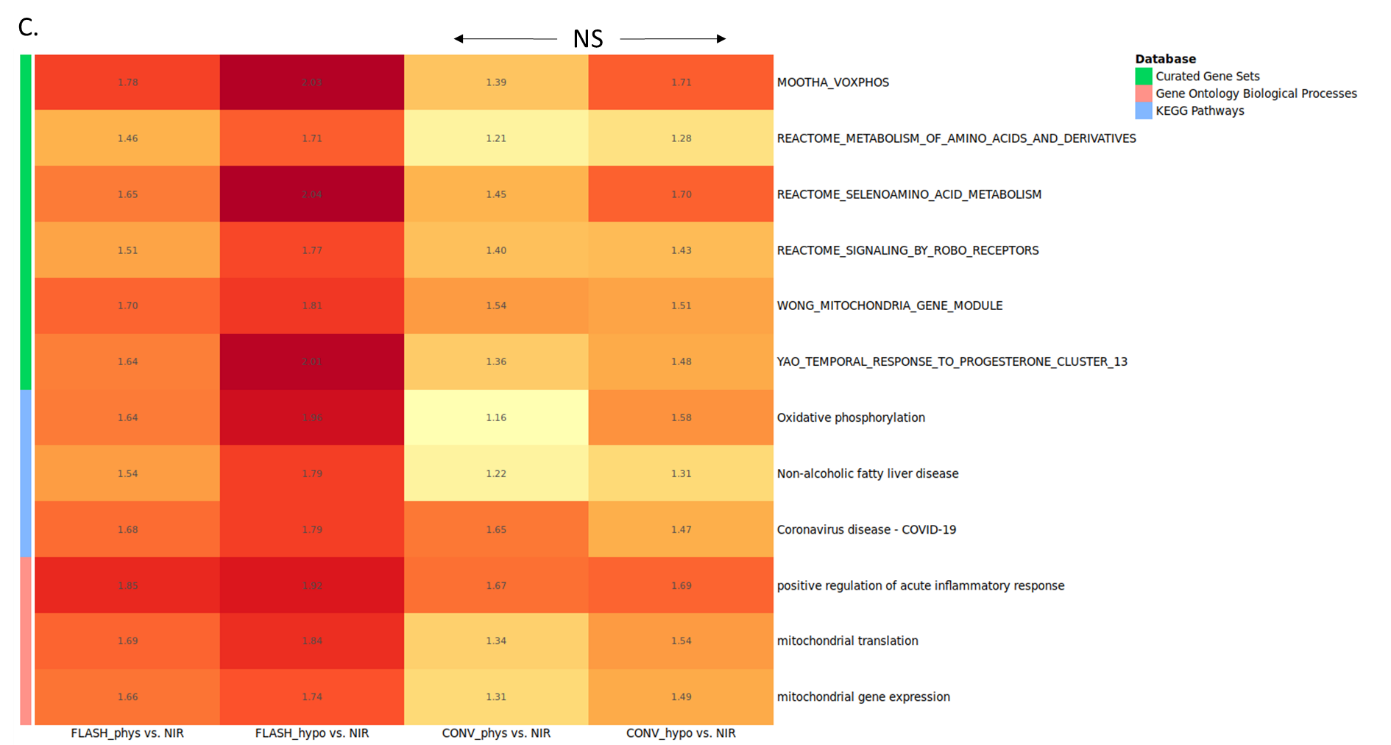

**Fig. S6.** Comparison overlap analysis of GSEA at one week post-recurrence. (A) Principal components analysis (PCA) performed to visualize sample-to-sample distances based on the top two principal components (PC1 and PC2), specifically for the physioxia and hypoxia conditions. (B) Heatmap generated with the same samples showing the top 500 genes – columns (samples) and rows (genes, Z-score) were clustered hierarchically. (C) Gene set enrichment analysis (GSEA) performed at the late time point comparing each treatment group with the non-irradiated (NIR) controls. This figure displays the FLASH-RT-specific gene set enrichments upregulated at the late time point. Numbers shown represent the normalized enrichment scores calculated by GSEA. Only “FLASH_phys vs. NIR” and “FLASH_hypo vs. NIR” comparisons statistically significant (FDR-adjusted p-value < 0.05); NS, not significant.

**Supplemental References**

1.	Jaccard M, Durán MT, Petersson K, Germond J-F, Liger P, Vozenin M-C, et al. High dose-per-pulse electron beam dosimetry: Commissioning of the Oriatron eRT6 prototype linear accelerator for preclinical use. Medical Physics. 2018;45:863–74.

2.	Liao Y, Shi W. Read trimming is not required for mapping and quantification of RNA-seq reads at the gene level. NAR Genomics and Bioinformatics. 2020;2:lqaa068.

3.	Dobin A, Davis CA, Schlesinger F, Drenkow J, Zaleski C, Jha S, et al. STAR: ultrafast universal RNA-seq aligner. Bioinformatics. 2013;29:15–21.

4.	Li H, Handsaker B, Wysoker A, Fennell T, Ruan J, Homer N, et al. The Sequence Alignment/Map format and SAMtools. Bioinformatics. 2009;25:2078–9.

5.	Kluin RJC, Kemper K, Kuilman T, de Ruiter JR, Iyer V, Forment JV, et al. XenofilteR: computational deconvolution of mouse and human reads in tumor xenograft sequence data. BMC Bioinformatics. 2018;19:366.

6.	R Core Team. R: A Language and Environment for Statistical Computing. R Foundation for Statistical Computing [Internet]. 2021; Available from: https://www.R-project.org/

7.	RStudio Team. RStudio: Integrated Development Environment for R. RStudio, PBC [Internet]. 2021; Available from: http://www.rstudio.com/

8.	Liao Y, Smyth GK, Shi W. featureCounts: an efficient general purpose program for assigning sequence reads to genomic features. Bioinformatics. 2014;30:923–30.

9.	Love MI, Huber W, Anders S. Moderated estimation of fold change and dispersion for RNA-seq data with DESeq2. Genome Biology. 2014;15:550.

10.	Wu T, Hu E, Xu S, Chen M, Guo P, Dai Z, et al. clusterProfiler 4.0: A universal enrichment tool for interpreting omics data. The Innovation. 2021;2:100141.

11.	Wickham H. ggplot2: Elegant Graphics for Data Analysis [Internet]. Springer-Verlag New York; 2016. Available from: https://ggplot2.tidyverse.org

12.	Kolde R. pheatmap: Pretty Heatmaps. R package version 1.0. 12. CRAN R-project org/package= pheatmap. 2019;

13.	Luo W, Brouwer C. Pathview: an R/Bioconductor package for pathway-based data integration and visualization. Bioinformatics. 2013;29:1830–1.

14.	Benjamini Y, Hochberg Y. Controlling the False Discovery Rate: A Practical and Powerful Approach to Multiple Testing. Journal of the Royal Statistical Society: Series B (Methodological). 1995;57:289–300.
